# GENE FLOW AND CONSERVATION GENOMICS OF *DUDLEYA ABRAMSII* SSP. *SETCHELLII*, AN ENDANGERED SUCCULENT IN SANTA CLARA COUNTY

**DOI:** 10.64898/2026.11.09.764340

**Authors:** Dante Cable, Karina Martinez, Evan Hackstadt, Victor Rossi, Abigail Sauter, Isabella Fasolino, Melissa Lopez, Charlotte Ryder, Christal Niederer, Jimmy Quenelle, Christopher Schwind, Marissa Kent, Stuart B. Weiss, Chris Baysdorfer, Weilun Tan, Lan Wei, Stephen Ward Mccabe, Amanda Hoppers, Scott Hansen, Bryan Bach, Shana L. Mcdevitt, Carrie Rose Miller, Justen B. Whittall

## Abstract

Understanding how gene flow shapes the distribution of genetic variation is essential for the conservation of rare and geographically restricted species. We investigated genomic structure, clonality, and mechanisms of gene flow in the federally endangered, serpentine endemic *Dudleya abramsii* ssp. *setchellii* (Jeps.) Moran using low-coverage whole-genome resequencing of 52 individuals representing 13 populations of *D. a.* ssp. *setchellii*, four populations of the geographically adjacent *D. cymosa* (Lem.) Britton & Rose, and two other subspecies of *D. abramsii* from San Luis Obispo County. Genome-wide comparisons were combined with analyses of clonality, isolation by distance, principal component analysis, phylogenetic reconstruction, pollinator observations, and a controlled seed dispersal experiment. Strong isolation by distance was detected both within a focal population and among populations across the species’ range, indicating that genetic differentiation increases predictably with geographic separation. Clonal reproduction was present, but restricted to individuals separated by only a few centimeters, contributing little to broader spatial genetic structure. Genomic analyses recovered *D. a.* ssp. *setchellii* as a distinct lineage exhibiting geographic subdivision within the species and identified a putative hybrid with *D. cymosa*. Field observations identified bumblebees as the dominant floral visitors, suggesting the potential for pollen-mediated gene flow, whereas a controlled seed dispersal experiment showed that most seeds traveled only a few meters under the artifically optimal experimental wind conditions. Together, these results suggest that limited seed dispersal may contribute to the pronounced spatial genetic structure observed in this serpentine endemic, although the relative contributions of pollen-mediated gene flow, geographic isolation, and genetic drift remain unresolved. These findings provide a genomic framework for conservation and identify geographically structured populations (east vs. west of Coyote Valley) as important units for long-term management.

---

Understanding how gene flow shapes the distribution of genetic variation is central to both evolutionary biology and the conservation of rare species. In plants, gene flow occurs through two primary mechanisms, pollen movement and seed dispersal, which can operate over different spatial scales. Differences in the distances traveled by pollen and seeds, together with geographic isolation and genetic drift, shape the spatial distribution of genetic variation within and among populations. Identifying genetically distinct populations can therefore guide conservation strategies aimed at preserving genetic diversity across a species’ range. Genetic diversity provides the raw material for adaptive evolution, while restricted gene flow and genetic drift can erode this variation, potentially limiting the ability of small, isolated populations to respond to environmental change.

When dispersal is spatially restricted, populations are expected to exhibit isolation by distance (IBD), in which individuals that are geographically closer are more genetically similar than those farther apart (Wright 1943; Slatkin 1993; Rousset 1997; Perez et al. 2018). This pattern reflects the cumulative effects of limited gene flow and can arise at multiple spatial scales, from fine-scale structure within populations to broader patterns across a species’ range as demonstrated in seep monkeyflowers (*Mimulus guttatus*) in western North America (Twyford et al. 2020). Understanding the scale and strength of IBD is particularly important in conservation contexts, as it can help identify populations that are genetically isolated or that harbor unique genetic variation.

Clonal reproduction can further complicate the interpretation of spatial genetic structure. In species capable of vegetative propagation, clusters of genetically identical individuals may arise from a single parent, potentially inflating local genetic similarity and mimicking patterns expected under limited dispersal. Clone size and number of ramets can vary widely, as in the rare sedge from Australia, *Lepidosperma* sp. (Halkett et al. 2005), where extensive clonal reproduction has substantially reduced the number of genetically unique individuals (Binks et al. 2015). Clonal propagation provides reproductive assurance in small, isolated populations like the insular *Ruta microcarpa* from the Canary Islands, where it essentially guarantees reproductive success even under harsh environmental stress (Meloni et al. 2013). Here, clonal propagation may benefit rare species in the short-term, but limit genetic recombination and evolutionary potential in the long term. In addition, clonal reproduction can decouple census population size from effective population size, affecting the maintenance of genetic diversity and long-term population viability. Determining the extent and spatial scale of clonality is therefore essential for accurately interpreting spatial genetic patterns and for informing conservation strategies. However, identifying clones from genomic data requires distinguishing technical variation among samples from genuine genetic differences among individuals. One approach is to generate artificial clones by independently processing replicate samples from the same individual, thereby establishing an empirical threshold of genetic differentiation below which individuals can be considered putative clones.

In order to investigate IBD and the effects of clonality on the distribution of genetic diversity, we focused on serpentine habitats whose insular nature provides a powerful natural framework. Characterized by soils with low Ca:Mg ratios, excess heavy metals, and a generally nutrient-poor composition, serpentine substrates support numerous edaphic endemics (255 species, approximately 14% of the California Flora) that are often restricted to discrete outcrops embedded within a matrix of this otherwise unsuitable habitat (Safford and Miller 2020). These outcrops function as terrestrial analogs of islands, where isolation among serpentine patches can limit both pollen- and seed-mediated gene flow. Such fragmentation is expected to generate strong spatial genetic structure and pronounced isolation by distance (Wolf et al. 2000; Harrison and Inouye 2002; Anacker et al. 2011).

*Dudleya abramsii* ssp. *setchellii* (Jeps.) Moran (Santa Clara Valley Dudleya,) is a federally endangered perennial succulent endemic to serpentine outcrops in southern Santa Clara County, California (United States Fish and Wildlife Service 2021) (Fig. 1). The species occupies rocky outcrops amidst serpentine grassland with populations often separated by unsuitable habitat (serpentine grassland, non-serpentine habitat, and increasingly urban development). These characteristics make *D. a.* ssp. *setchellii* an ideal species for examining how gene flow operates across both naturally and anthropogenically fragmented landscapes. For pollen-mediated gene flow, the species produces relatively large floral displays with 10-25 flowers per inflorescence and 1-7 inflorescences per rosette. Flowers produce abundant pollen, but little to no nectar reward, consistent with pollen-foraging insect pollinators. It is capable of autogamous self-pollination in a pollinator free greenhouse (Whittall unpublished data), but the degree of outcrossing is unknown. Regarding seed dispersal, seeds are extremely small (<1 mm), lack specialized structures for dispersal, and are likely dispersed passively by wind as suggested by similarities to other members of the genus (Jepson Flora Project 2026). Together, these traits suggest that pollen and seed movement may operate over small spatial scales, although the relative contributions of these processes to gene flow in *D. a.* ssp. *setchellii* remain unknown.

**FIG. 1.**
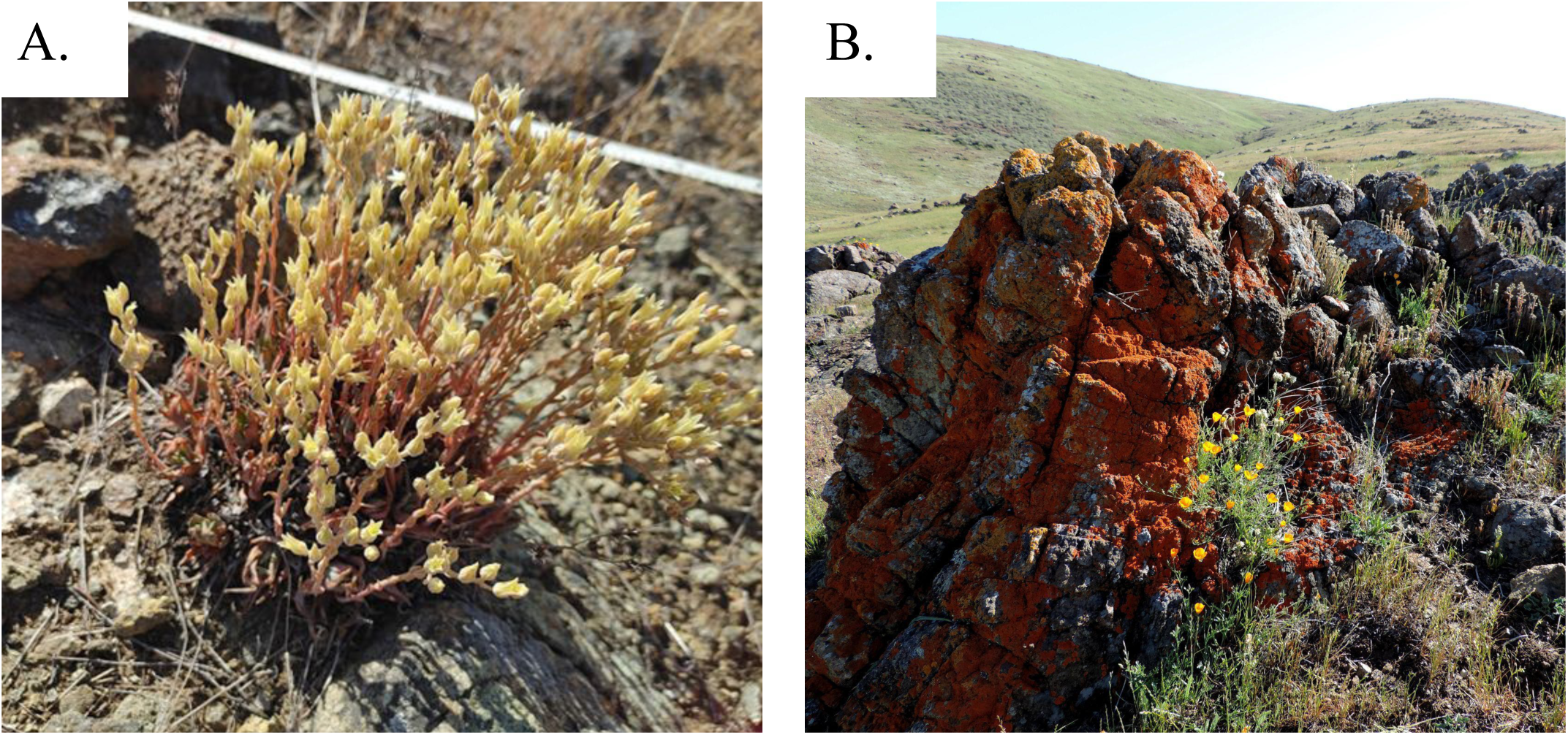
*Dudleya abramsii* ssp. *setchellii* is a serpentine endemic succulent. (A) A profusely flowering plant with multiple adjacent rosettes at “Dude Rock” along Coyote Ridge. (B) *D. a.* ssp. *setchellii* is restricted to serpentine outcrops that function as island-like habitats.

In addition to within- and among-population processes, interspecific gene flow may shape genetic structure in this system. *Dudleya abramsii* ssp. *setchellii* co-occurs with the widespread *D. cymosa* on adjacent serpentine and non-serpentine rocky outcrops, respectively (Jepson Flora Project 2026). The taxa differ in leaf morphology, inflorescence architecture, flower color, and summer dormancy (*D. a.* ssp. *setchellii* folds its drying leaves inward around the growing rosette in summer and fall), but share a diploid chromosome number (2n = 34) and overlapping spring flowering periods, providing opportunities for interspecific pollen transfer, particularly if they share pollinators. Although hybridization has not been documented between these taxa, their geographic proximity, reproductive overlap, and close evolutionary relationship (Yost et al. 2013) suggest that introgression is possible. Gene flow from the widespread *D. cymosa* could threaten the rare endemic through genetic assimilation and loss of locally adapted variation (Ellstrand and Elam 1993; Levin et al. 1996; Rhymer and Simberloff 1996), or alternatively, increase genetic diversity and evolutionary potential. Regardless of its evolutionary consequences, recent hybridization may produce individuals with intermediate genomic variation and morphological traits (e.g., leaf shape) when grown in a common garden.

Despite its conservation importance, genomic resources for *D. a.* ssp. *setchellii* remain limited. Currently, the only publicly available *Dudleya* nuclear genome assembly is for *D. densiflora*, with low contiguity (281 Mb across 162,434 contigs), along with a chloroplast genome for *D. farinosa* (NC_085682). Although recent population genomic studies have begun to characterize other *Dudleya* species using ddRAD-Seq (Iacuaniello et al. 2025), previous investigations of relationships involving *D. a.* ssp. *setchellii* have relied primarily on morphology and a limited number of molecular markers (Yost et al. 2013; Guilliams and Hasenstab-Lehman 2021), which lack the resolution needed to investigate fine-scale genetic structure, clonality, and introgression. The diploid chromosome number shared by *D. a.* ssp. *setchellii* and its close relatives (n = 17) facilitates the development of genome-scale resources (Jepson Flora Project 2026).

Previous phylogenetic analyses using nuclear ITS and chloroplast markers place *D. a.* ssp. *setchellii* in the *Virens* clade alongside two subspecies of the widespread, morphologically variable *D. cymosa* (*D. c.* ssp. *cymosa* and *D. c.* ssp. *paniculata*) and other subspecies of *D. abramsii* (Yost et al. 2013; McCabe 2012, 2024). Yost et al. (2013) also suggested the possibility of gene flow among these taxa where their distributions overlap, including in Santa Clara County. These relationships have contributed to a complex taxonomic history: *D. a.* ssp. *setchellii* has been recognized as a subspecies of *D. cymosa* (Moran 1951), a distinct species (Britton and Rose 1903; Moran 1957), and currently as a subspecies of *D. abramsii*. Two other serpentine-endemic subspecies of *D. abramsii* (ssp. *murina* and ssp. *bettinae*) occur more than 200 km to the south in San Luis Obispo County and share the unusual characteristic of leaf bases turning red when detached from the rosette (McCabe 2024). Whether the genetic similarities among these taxa reflect shared ancestry, recent divergence, or ongoing gene flow remains unresolved.

In this study, we integrate genomic comparisons with field and greenhouse studies to investigate the mechanisms and consequences of gene flow in *D. a.* ssp. *setchellii*. Specifically, we address gene flow at three hierarchical levels. First, we quantify the extent of clonal reproduction and test for isolation by distance within one focal population to assess fine-scale spatial genetic structure. Second, we evaluate isolation by distance among populations across the species’ range to determine how geographic isolation shapes regional genetic connectivity. Third, we examine evolutionary relationships among *D. a.* ssp. *setchellii* and closely related taxa to assess evidence for hybridization and introgression. We then interpret these genomic patterns in the context of pollinator observations and seed dispersal estimates to evaluate the mechanisms underlying the detected patterns of gene flow. By linking genetic structure across spatial scales to the biological processes of pollen and seed movement, this study provides insight into the factors that shape genetic diversity in a fragmented serpentine endemic and informs conservation strategies for maintaining its evolutionary potential.

## Methods

### Study System and Sampling

To characterize genetic structure within and among populations, we collected leaf tissue of *D. a. ssp. setchellii* from serpentine outcrops in southern Santa Clara County, California. Regional sampling included 13 populations spanning the known range of *D. a.* ssp. *setchellii*, four geographically adjacent populations of *D. cymosa*, and samples from two serpentine endemic subspecies of *D. abramsii* from San Luis Obispo County for broader phylogenetic context (Fig. 2, Table S1). At each site, 3–5 young leaves were collected from the central rosette of each plant and stored at 4°C until they were transferred to a −20°C freezer prior to DNA extraction.

**FIG. 2.**
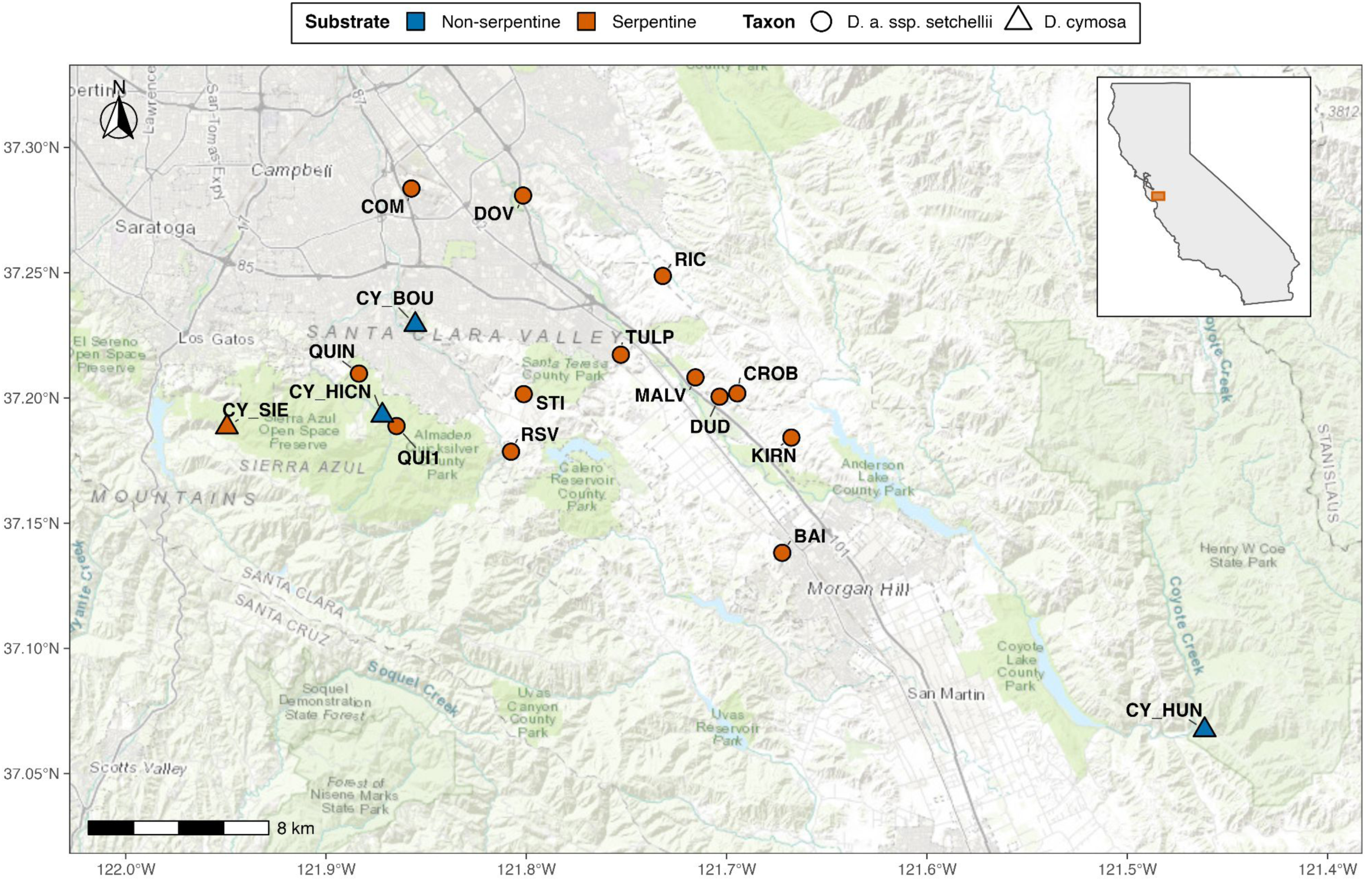
The approximate locations of sampled *Dudleya abramsii* ssp. *setchellii* populations in southern Santa Clara County, California (inset) are indicated by circles. Circles denote *D. a.* ssp. *setchellii* populations and triangles denote *D. cymosa* populations. Samples from two other subspecies of *D. abramsii* from San Luis Obispo County are not shown. Population abbreviations beginning with "CY" identify *D. cymosa* populations; all other population codes identify *D. a.* ssp. *setchellii* populations (see Table S1 for locality information). Population symbols are colored by substrate: orange = serpentine and blue = non-serpentine.

To assess fine-scale clonality and isolation by distance within a single focal population, we intensively sampled *D. a.* ssp. *setchellii* at Almaden Quicksilver County Park. We collected leaves at varying distances along five 50-100m transects (distances ranged from 1cm to 79.2m based on plant availability; Fig. S1). Leaves were intentionally collected from immediately adjacent plants (<10cm) and then from more geographically distant plants (up to ∼80m). In total, leaf tissue from 40 plants was collected (eight plants per transect x five transects) while also recording physical distances in cm among sampled individuals so that genetic and geographic distance could be compared directly. After DNA extraction, library preparation and sequencing, 32 individual samples with the highest DNA extraction concentrations and optimal library preps based on fragmentation and yield were sequenced for the within-population genomic analyses.

### Genome Size Estimation

To confirm the ploidy level and determine the amount of sequencing required, genome size was estimated by flow cytometry. Fresh leaf tissue from *D. a.* ssp. *setchellii* was analyzed independently at the Green Biome Institute (California State University, East Bay) and by Lifeasible (Shirley, NY). The CSU East Bay analysis focused on a single individual (see “Dude Rock” below) and used fava bean and tomato as a size standard, whereas the Lifeasible analysis used diploid *Hemerocallis* cultivars (∼8.99 pg 2C DNA) and included five *D. a.* ssp. *setchellii* samples.

Reference Genome

A reference genome for *D. a.* ssp. *setchellii* was generated from a single individual collected from “Dude Rock”, a nickname for a particularly dense cluster of plants on Máyyan ‘Ooyákma Coyote Ridge Open Space Preserve (Table S1). The reference genome required unfragmented DNA, longer reads and higher sequencing coverage than the population resequencing dataset. In collaboration with the Green Biome Institute at California State University, East Bay, the reference genome methods are described below.

*Reference Genome DNA Extraction –* DNA extraction was conducted using a modified CTAB-based protocol adapted from Aboul-Maaty and Oraby (2019). Approximately 300 mg of leaf tissue was homogenized with a mortar and pestle under liquid nitrogen and extracted in 1 mL of 2% CTAB buffer supplemented with 10 µL of 2-mercaptoethanol. Samples were incubated at 65°C for 1 h with agitation (300 rpm) and periodic inversion, followed by two chloroform alcohol (24:1) extractions. DNA was precipitated by adding 300 µL of 5 M NaCl, 60 µL of 3 M sodium acetate, and 600 µL of ice-cold isopropanol to 600 µL of aqueous phase. To minimize DNA shearing, solutions were gently mixed and transferred using wide-bore pipette tips into DNA LoBind tubes. DNA pellets were washed twice with 80% ethanol and resuspended in 35 µL of nuclease-free water. The extracted DNA was assessed for purity using a NanoDrop spectrophotometer, for fragment size distribution, and concentration using the Agilent TapeStation 4200 with the Genomic DNA ScreenTape assay, and for DNA concentration using a Qubit 4 Fluorometer with the dsDNA High Sensitivity assay kit.

*Reference Genome Library Preparation and Sequencing* – We prepared a high-molecular-weight genomic DNA library from 2 µg of DNA using the Oxford Nanopore Ligation Sequencing Kit V14 (SQK-LSK114), following the manufacturer’s protocol with modifications to maximize recovery of long DNA fragments. We extended the DNA repair and end-preparation incubations to 30 min each at 20°C and 65°C, AMPure XP bead binding and DNA elution to 15 min each, and adapter ligation to 30 min at 23°C. We used NEBNext Quick T4 DNA Ligase for adapter ligation and minimized DNA shearing through gentle mixing, wide-bore pipette tips, and DNA LoBind tubes. The library was purified using AMPure XP beads, and fragment size distribution and concentration were assessed using an Agilent TapeStation 4200 with the Genomic DNA ScreenTape assay and a Qubit 4 Fluorometer with the dsDNA High Sensitivity assay, respectively. We sequenced approximately 30 fmol of the prepared library on an Oxford Nanopore PromethION 2 Solo using an R10.4.1 flow cell for approximately 48 h, with real-time basecalling in MinKNOW using the super-accuracy (SUP) model.

*Reference Genome Assembly* – The assembly was performed on Amazon Web Services (AWS) using the Ubuntu 24.04 operating system. Low-quality reads with a Q score smaller than 10 were removed. De novo genome assembly was performed using hifiasm v0.25.0 with default parameters optimized for long-read assembly. To improve assembly quality, redundant haplotigs and duplicated contigs resulting from heterozygosity were removed using purge_dups v1.2.6.

Next, contigs classified as microbiome by kraken2 v2.1.6 in the draft assembly were removed. Furthermore, contigs containing mitochondrial and chloroplast-related sequences were identified using BLASTn and then separated into separate FASTA files. Lastly, the host assembly genome was assessed using QUAST v5.3.0 to calculate assembly metrics including total genome size, contig N50, number of contigs, and maximum contig length. Genome completeness was evaluated using BUSCO v5.6.1 using the Viridiplantae Odb10 lineage database.

### DNA Extraction, Library Preparation, and Resequencing

For population resequencing, total genomic DNA was extracted from frozen leaf tissue using the NucleoSpin Plant II kit (Macherey-Nagel, Düren, Germany) with lysis buffer PL1. Before extraction, the leaves’ glaucous waxy coating was removed with a kimwipe, 90–100 mg tissue was weighed, and leaves were chopped into fragments smaller than a 5 mm square. Tissue was homogenized in 2 mL screw-cap tubes containing zirconia/silica beads with a bead beater (BioSpec Products, Bartlesville, OK, USA) for 2 min, followed by extraction according to the manufacturer’s protocol. DNA was eluted in 100 µL pre-warmed PE buffer and stored at −20°C. Extract concentration and quality were assessed using a Qubit 3.0 Fluorometer with dsDNA High Sensitivity Assay Kit (Thermo Fisher Scientific, Waltham, MA, USA), Nanodrop (Thermo Fisher Scientific, Waltham, MA, USA), and agarose gel electrophoresis.

Genomic libraries for Illumina resequencing were prepared using the NEBNext Ultra II Express FS DNA Library Prep (New England Biolabs, Ipswich, MA) workflow with slight modifications to the manufacturer’s protocol. We optimized a miniaturized format (1/3x original reaction volume), with enzymatic fragmentation for 17 min targeting inserts of roughly 300–350 bp. Universal adaptors (NEB) and unique dual indexes (QB3, UC Berkeley, CA) were incorporated followed by a PCR step of six cycles to finish the libraries. Resulting libraries were washed twice with ethanol and adaptor dimers were removed using a 0.7× SPRI bead cleanup with a final elution in 30 µL. Library size distributions and concentrations were assessed on an Agilent TapeStation D1000 (Agilent Technologies, Santa Clara, CA, USA) and an additional round of bead cleanup was performed at 0.7x SPRI bead ratio to remove any residual adapter-dimers and primers. Finally, equimolar pools were assembled using concentration estimates from the desired fragment range (200-1,000 bp). Sequencing was performed at QB3 (Berkeley, CA) using Illumina paired-end 150 bp reads. Samples were sequenced with a target minimum of 5X coverage per sample on NovaSeq X 10B instrument (375 Gbp per run).

### Read Processing, Alignment, and Genome Comparisons

Raw paired-end reads (fastq) were filtered and trimmed with fastp v0.22.0 (Chen et al. 2018), then aligned to the *D. a.* ssp. *setchellii* reference genome generated herein using BWA-MEM v0.7.18 (Li 2013). Alignments were sorted and indexed with SAMtools v1.6 (Danecek et al. 2021) generating BAM files. Genotype likelihoods and allele frequency likelihoods were estimated using ANGSD v0.933 (Korneliussen et al. 2014) on all 52 individuals. Beagle-formatted genotype likelihoods (-doGlf 2) were generated using the SAMtools model (-GL 1). Major and minor alleles and their frequencies were inferred from the genotype data (-doMajorMinor 1, -doMaf 1). Hard genotype calls were stored in a BCF file (-doGeno 32, -doPost 1, -doBcf 1). The frequency of each base was counted (-doCounts 1) for a sequencing depth report (-doDepth 1, -maxDepth 1000). Only unique reads (-uniqueOnly 1) with a minimum mapping quality of 30 (-minMapQ 30) and sites with a mean base quality of 20 (-minQ 20) were considered. Five bases were trimmed from each end of reads (-trim 5). Notably, analysis was not restricted only to variable sites, allowing most of the genome (∼232 million sites) to be considered. Pairwise genetic distances were calculated with ngsDist v1.0.10 (Vieira et al. 2016) with the --avg_nuc_dist and --evol_model_0 flags to obtain raw p-distances, and the -- pairwise_del flag to maximize the number of sites kept for pairwise comparisons. All bioinformatic analyses were conducted using SCU’s high performance computing cluster (WAVE). An overview of the bioinformatics workflow is shown in Fig. S2 and read processing statistics are reported in Fig. S3 including the distribution of post-filtering coverage per sample.

### Clonality and Isolation by Distance in *D. a.* ssp. *setchellii*

To estimate clonality within a single focal population (Quicksilver North), we quantified the amount of sequence divergence introduced during sample processing and sequencing to determine a p-distance cutoff for assessing clonality. To do this, nine artificial clone pairs were generated by taking paired tissue samples from the same individual and processing them independently from DNA extraction through sequence analysis (Fig. S4). The distribution of genetic distances among these artificial clone pairs was then used to establish a threshold for identifying putative clones in the field-collected dataset. Pairwise genetic distances among all samples were compared with pairwise geographic distances measured during field sampling to determine the geographic distance between putative clones.

Isolation by distance in *D. a.* ssp. *setchellii* was evaluated at two spatial scales: (1) within the focal Quicksilver North population and (2) among populations spanning the species’ geographic range. Artificial clone replicates used to calibrate the clonality threshold and putative clonal genets identified among field-collected samples were each arbitrarily reduced to a single representative sample prior to analysis to avoid pseudoreplication. For the among population isolation by distance analysis, the numerous QUIN samples were averaged into a single representative sample for that population. Genetic (p-distance) and geographic distance matrices were compared using Mantel tests with 9,999 permutations. Analyses were conducted using both raw geographic distances and log₁₀-transformed geographic distances (log₁₀[distance + 1]) to evaluate whether a logarithmic transformation better described the relationship between geographic and genetic distance, as expected if genetic differentiation accumulates more rapidly over short than long distances. Genetic and geographic distance matrices and associated R script are available from Figshare (Supplemental Materials).

### PCA and Phylogenetic Analyses Including Outgroups

To visualize genomic relationships within *D. a.* ssp. *setchellii* and among it and its closest relatives, we performed principal component analysis (PCA) and phylogenetic analyses using genotype likelihoods generated with ANGSD from low-coverage whole-genome resequencing data (described above). PCA was performed with PCAngsd v1.36.4 (Meisner and Albrechtsen 2018) and the first two principal component axes were used to visualize genomic similarities among the samples. Phylogenetic relationships were inferred using two complementary approaches to assess the consistency of inferred relationships: neighbor-joining based on pairwise genetic (p-) distances and maximum likelihood implemented in IQ-TREE using a custom neighbor-joining script (“matrix2tree” available in the Supplemental materials) and IQ-TREE v3.0.1 (Wong et al. 2026). Maximum-likelihood analyses were performed using the GTR+ASC substitution model, with ascertainment-bias correction to account for the exclusion of invariant sites from the SNP-only alignment. We selected GTR as a flexible, general nucleotide substitution model that allows unequal substitution rates and nucleotide frequencies. Phylogenetic trees were visualized in Geneious Prime v.2026.1.2 (Biomatters, Auckland, New Zealand).

Branch support was assessed using 1,000 ultrafast bootstrap replicates. To determine whether genomes of *D. a.* ssp. *setchellii* are more similar to *D. cymosa* or the two other sampled subspecies of *D. abramsii,* we compared mean p-distances and tested these for significance using a permutation test with 1,000 replicates where taxon labels were permuted at the individual level to avoid pseudoreplication associated with pairwise distance calculations.

### Leaf Morphometrics

To compare leaf dimensions among *D. a*. ssp. *setchellii* and its putative close relatives (*D. cymosa* and other serpentine endemic subspecies of *D. abramsii*) field-collected plants were grown for more than one year in a common garden at the SCU Greenhouse Facility. Samples were collected from the field under our USFWS LAG permit, then planted in 1 gallon pots using standard potting soil (Miracle Grow Potting Mix). The greenhouse temperature was maintained between 60-90 F with a swamp cooler running in the heat of summer. Supplemental crop lights ran for 14hrs/day throughout the year. Plants were watered 1-3x per week equally and were rotated arbitrarily every 2-3 weeks.

For each plant, the three longest leaves were measured with digital calipers. Leaf length was measured from the point of attachment at the rosette to the leaf tip, and width was measured at the base, midpoint, and widest point. Measurements were averaged across the three leaves to obtain one plant-level value for each trait. Because trait distributions did not meet parametric assumptions and some groups had small sample sizes, differences among groups were evaluated using Kruskal–Wallis tests followed by Dunn pairwise comparisons with Holm adjustment. Plants from Boulder Ridge were analyzed as a separate group (BOU; n = 4) because this population included CY_BOU, the genetically intermediate genomic sample that we regard as a putative hybrid (see Results below).

### Pollinator Observations

In order to characterize the potential for pollen-mediated gene flow, we recorded pollinator visitation at *D. a.* ssp. *setchellii* populations on serpentine outcrops in southern Santa Clara County during peak flowering season in June 2024. Ten digital video camcorders (Canon VIXIA HF series; 8 GB internal memory) were deployed simultaneously across two study sites on 13 and 18 June 2024 (Fig. S5). Cameras were mounted on tripods approximately 20–40 cm from focal plants and recorded from shortly after sunrise until early evening. Temperatures were typical for this region at this time of year ranging from ∼15° C in the morning to a high of 21.1° C and 27.2° C in the early afternoons on the two days we monitored pollinators. Each camera monitored one to four flowering individuals bearing approximately 15–40 open flowers (Table S2).

All videos were reviewed manually to quantify pollinator visitation. A visit was recorded when an insect contacted the reproductive structures of a single flower (even though most recordings had multiple flowers per frame). Both the number and duration of every flower visited were recorded. Visit duration was defined as the elapsed time from first to last floral contact. Pollinators were identified to major taxonomic groups (pollinator guilds) based on visible morphological characteristics and comparison with a reference insect collection assembled from the study sites. These observations were used to identify the primary pollinators and characterize visitation frequency and duration as indicators of the potential for pollen-mediated gene flow. This method quantifies pollinator visitation frequency (a visit that contacts reproductive structures) but does not address pollination effectiveness, pollen transfer or seed set.

### Seed Dispersal Experiment

To estimate the potential for wind-mediated seed dispersal under controlled conditions, seed dispersal trials were conducted indoors using field-collected *D. a.* ssp. *setchellii* seeds. Seed morphology was first documented under a stereomicroscope (AmScope SM-3TZZ-144A-5M3-B; 3.5–180× magnification) equipped with a 5 MP digital camera. Seed dispersal trials consisted of ten replicates of 100 seeds each. Seeds were released from a height corresponding to the average height of naturally dehiscing infructescences (30 cm) while exposed to airflow generated by a leaf blower positioned 1 m from the release point (Fig. 8B). Airflow at the seed release point was set to 33.5 mph, corresponding to the higher end of wind speeds recorded along Coyote Ridge using a Kestrel 5000 LiNK anemometer (Nielsen-Kellerman). The wind rose from Reid-Hillview airport (∼15km north of the center of *D. a.* ssp. *setchellii* geographic distribution) shows that wind speeds greater than 20 mph occur ∼0.5% of the time between August-November when *D. a.* ssp. *setchellii* fruits are dehiscing. In our experimental setting, seeds were dispersed onto a hallway floor covered with white butcher paper, and dispersal distances were measured in cm from the seed release point. Approximately 85% of released seeds were recovered and included in subsequent analyses.

The experiment was designed to estimate the wind-mediated seed dispersal under simplified, standardized conditions rather than to replicate natural field conditions. In contrast to the field, airflow generated by the leaf blower attenuated rapidly with increasing distance from the seed source, so air velocity was measured at increasing distances from the seed release point (Table S3). This experiment did not incorporate natural topography, vegetation, or turbulent wind patterns.

## Results

### Genome size

Mean flow-cytometry estimates were very similar among the *D. a.* ssp. *setchellii* samples and between the two labs (∼0.60 pg/2C, corresponding to about 588.8 Mb for the diploid genome) (Table S4). In comparison to a broader sampling of the genus including several samples with known chromosome counts (McCabe unpublished data), these estimates confirmed that all *D. a.* ssp. *setchellii* samples were diploid.

### Genome assembly

We combined short, paired-end reads from the Illumina platform (150bp x 2) with long reads on the Oxford Nanopore platform (>100,000bp/read) to assemble a reference genome with ∼100x coverage. The draft *D. a.* ssp. *setchellii* reference genome consists of 17 contigs with a total assembly length of 247 million bp (GenBank Genome Accession XXXXX) which is similar (but 16% smaller) to our genome size estimate from flow cytometry (294 million bp haploid estimate based on 0.60pg/2C) indicating potentially 47 million bp missing. These 17 contigs may represent the 17 unique chromosomes of *D. a.* ssp. *setchellii* which has 2n=34 (McCabe 2024). This reference genome was used for all downstream mapping and population genomic analyses. Although we also assembled complete cpDNA and mtDNA organellar genomes, we have not included them in these analyses. Furthermore, we have not intentionally masked any repetitive regions.

### Clonality

To assess the clonal capability of *D. a.* ssp. *setchellii*, we created nine artificial clones to quantify a p-distance genomic cutoff for assessing clonality. The mean genetic distance for the artificial clones was 0.04290 (when applied, this mean yields a conservative degree of clonality) with a maximum clonal cutoff value of 0.04773 (using the maximum value provides a more liberal interpretation of clonality, but may increase false positives - concluding a clone when really just genetically very similar, separate individuals). Teasing apart clones from very close relatives (i.e. self progeny) could be challenging if this species was an obligate selfer and therefore mostly homozygous. However, the genetic variation we see within populations (i.e. IBD within Quicksilver) indicates *D. a.* ssp. *setchellii* has a mixed mating system with modest genetic variation detected both within and among populations.

Using the mean artificial clone genetic distance cutoff, there were two separate pairs of field-collected plants that could be considered clones. The individuals in these two pairs were collected 3 cm and 6 cm apart from one another (Table 1). Broadening the clone cutoff by using the maximum genetic distance among artificial clones adds three more potential clones to this list. These three pairs were separated by 4 cm, 4 cm and 8 cm. Overall, the mean geographic separation between the five putative clones was 5 cm and four of the five putative clonal pairs were very close to the maximum cutoff (p-distance within 0.00551).

All putative clone pairs occurred within 8 cm. Among 11 sampled pairs of neighboring rosettes separated by ≤10 cm, five (45.5%) fell below the maximum artificial-clone threshold, whereas two (18.2%) fell below the mean threshold.

### Isolation by Distance

Significant isolation by distance was detected both within the focal Quicksilver North population (*r* = 0.490) and among populations (*r* = 0.656) using raw geographic distances (Mantel tests, *P* = 0.0001; Fig. 3, Fig. S6). The log₁₀ transformation of geographic distance strengthened the isolation-by-distance relationship at both spatial scales, increasing Mantel *r* from 0.490 to 0.524 within Quicksilver North and from 0.656 to 0.715 among populations (Table S5).

**FIG. 3.**
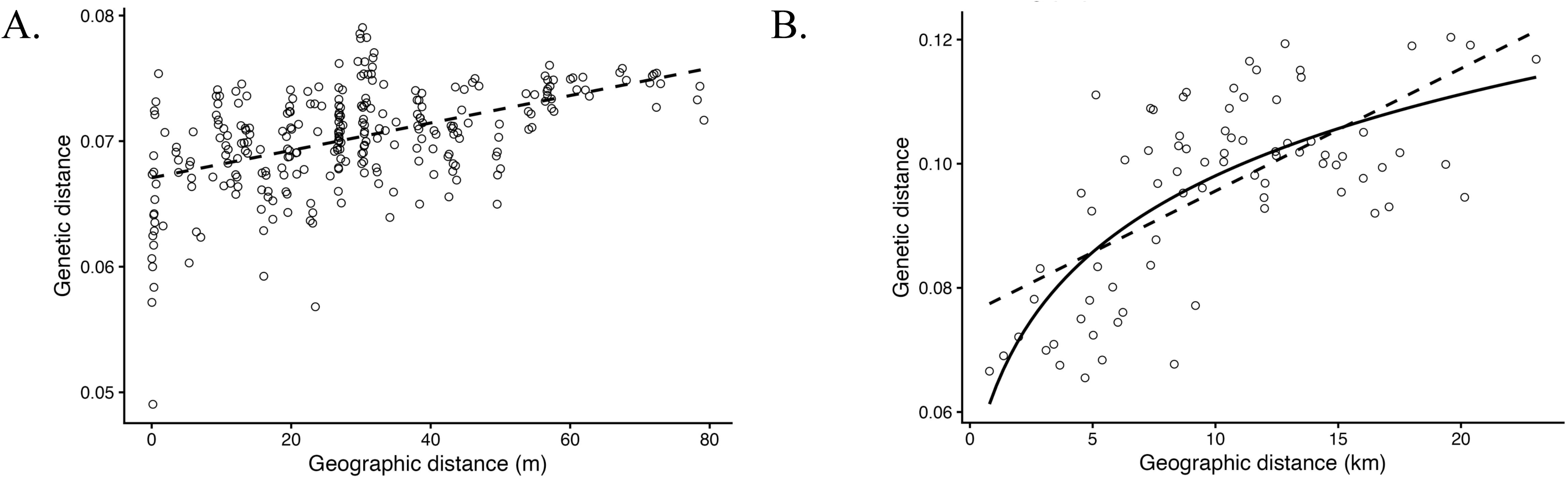
Genomic isolation by distance within and among *Dudleya abramsii* ssp. *setchellii* populations. (A) Relationship between pairwise genetic and geographic distance within the Quicksilver North population after retaining a single representative from each clonal genet (n = 24 individuals; 276 pairwise comparisons; Mantel r = 0.490; two-sided *P* = 0.0001). (B) Relationship between mean pairwise genetic and geographic distance among sampled *D. a.* ssp. *setchellii* populations (n = 13 populations; 78 pairwise comparisons; Mantel r = 0.656; two-sided *P* = 0.0001). Dashed lines show least-squares linear regressions using untransformed geographic distance in both panels. In panel B, the solid curve shows the regression fitted using log₁₀(distance + 1), displayed on the original geographic-distance scale. Individual points in (B) are labeled by their population names in Fig. S6.

### PCA

Principal component analysis of genomic diversity in *D. a.* ssp. *setchellii* (n = 45), *D. cymosa* (n = 4), and three samples representing two other subspecies of *D. abramsii* from San Luis Obispo County (n = 3) separated *D. a.* ssp. *setchellii* samples from the two comparison groups along PC1 which explained 31.64% of the variation (Fig. 4). One individual identified in the field as *D. cymosa* from Bould Ridge (CY_BOU) occupied an intermediate position between *D. cymosa* and *D. a.* ssp. *setchellii* along PC1. PC2 (9.37% of the variation) geographically separated populations within *D. a.* ssp. *setchellii*. All plants from Quicksilver North clustered together (n=33) and were situated more broadly adjacent to *D. a.* ssp. *setchellii* plants from the western portion of the species range (Fig. 4). PC3 (8.53%) clearly separated *D. cymosa* from the other subspecies of *D. abramsii* from San Luis Obsipo Co and PC4 (2.16%) further separated the *D. a.* ssp. *setchellii* samples (Fig. S7).

**FIG. 4.**
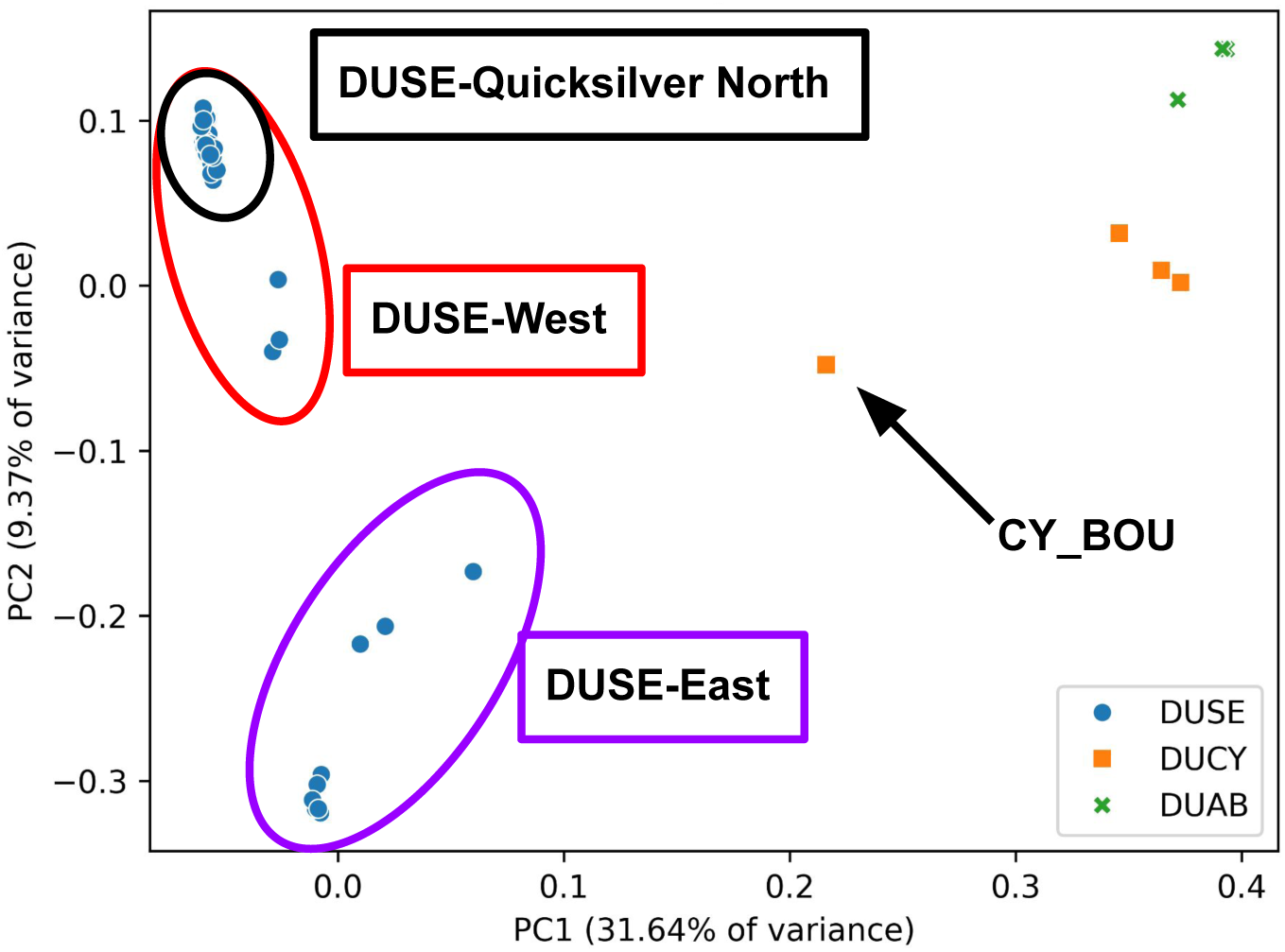
Principal component analysis based on genome-wide resequencing data from *D. abramsii* ssp. *setchellii* (n = 45; blue circles), *D. cymosa* (n = 4; orange squares), and three samples representing two other subspecies of *D. abramsii* from San Luis Obispo County (n = 3; green × symbols). Ellipses indicate geographic clustering within *D. a.* ssp. *setchellii* (black = Quicksilver North, red = western range, purple = eastern range). The arrow identifies CY_BOU, a genetically intermediate sample and therefore putative hybird between *D. a.* ssp. *setchellii* and *D. cymosa*. PC3 and PC4 are illustrated in Fig. S7.

### Phylogenetic Analysis

Neighbor-joining phylogenetic analysis using the genomic distance matrix (Fig. 5) recovered similar patterns identified in the PCA. The neighbor-joining tree shows a single grouping of all *D. a.* ssp. *setchellii* samples with geographically structured clusters therein. Further, it suggests that *D. a.* ssp. *setchellii* is more closely related to sampled populations of *D. cymosa* than it is to other sampled *D. abramsii* subspecies that are not identified as *D. a.* ssp. *setchellii*. The *D. cymosa* sample with an intermediate position in the PCA (CY_BOU), falls in between the *D. cymosa* and *D. a.* ssp. *setchellii* samples, closer to *D. a.* ssp. *setchellii* than to other sampled *D. cymosa* individuals. Broadly similar relationships were recovered in the maximum-likelihood analysis, where all but two nodes received 100% bootstrap support, although one *D. cymosa* sample from Hunting Hollow fell within the *D. a.* ssp. *setchellii* grouping (Fig. S8).

**FIG. 5.**
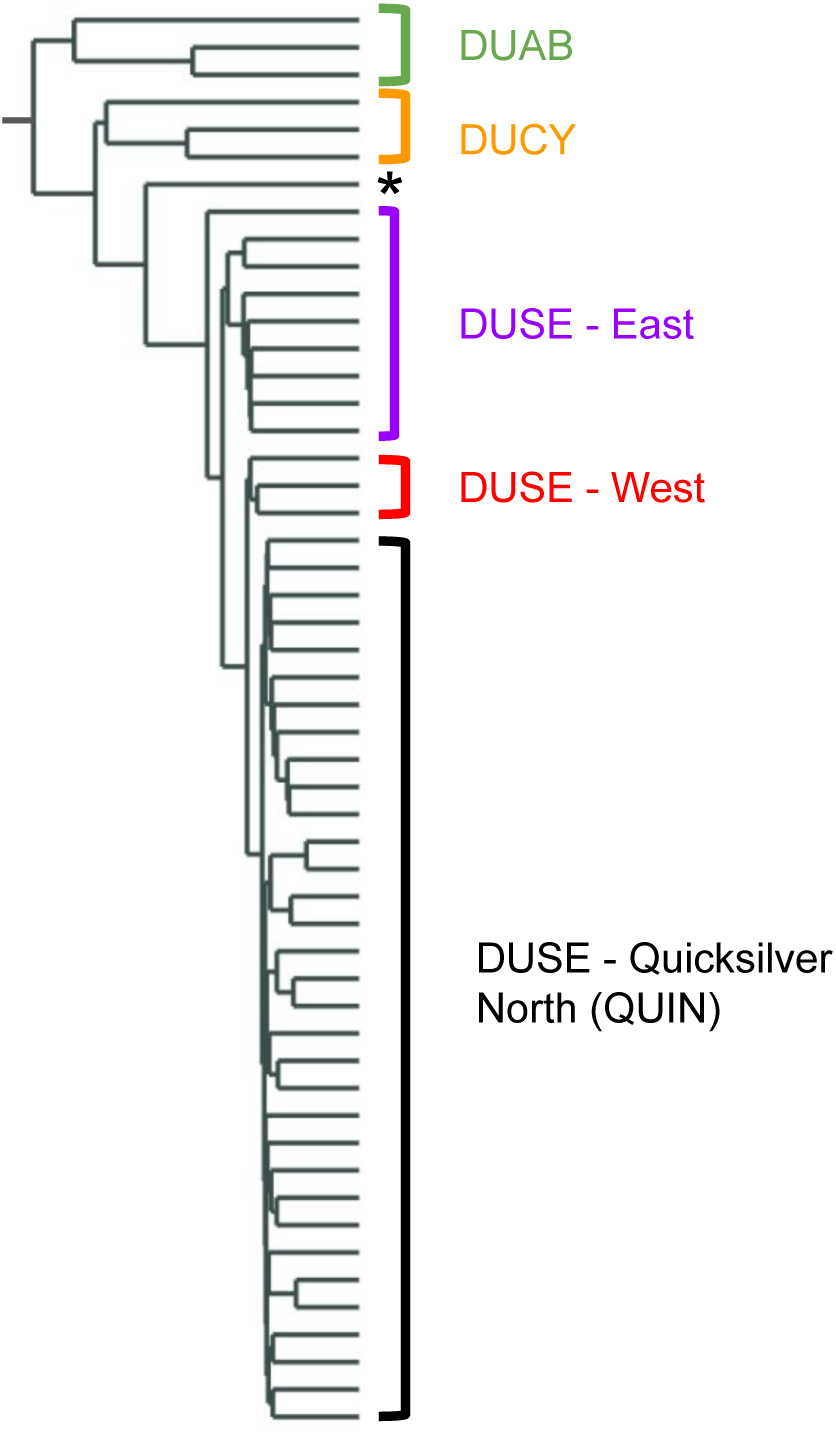
Neighbor-joining tree inferred from genome-wide pairwise p-distances among *D. abramsii* ssp. *setchellii*, *D. cymosa*, and three samples representing two other subspecies of *D. abramsii* from San Luis Obispo County. Brackets indicate groupings, including three geographically-defined groupings within *D. a.* ssp. *setchellii*: East, West (RSV, STI, QUI1), and Quicksilver North (QUIN). The black asterisk identifies CY_BOU (genetically intermediate sample and putative hybrid as shown in Fig. 4). “DUAB” denotes three samples representing two other subspecies of *D. abramsii* collected from San Luis Obispo County. Trees with complete sample names are included in Fig. S8.

Closer examination of the distance matrix indicates that *D. a.* ssp. *setchellii* is 56% more genetically similar to sampled populations of *D. cymosa* (N = 180 pairwise comparisons, mean p-distance = 0.016799, SE = 0.000153) than it is to the other two subspecies *D. abramsii* (N = 135 pairwise comparisons, mean = 0.026235, SE = 0.000283). In a permutation test, this genomic distance difference (-0.009436) is significant (*P* = 0.023).

The putative hybrid sample (CY_BOU) is nearly equidistant from *D. a.* ssp. *setchellii* and *D. cymosa*. It has mean genetic distances of 0.013352 to *D. a.* ssp. *setchellii* and 0.014949 to *D. cymosa*. These distances are only approximately half the distance from CY_BOU to the other two *D. abramsii* subspecies that were sampled from San Luis Obispo County (0.025393).

### Leaf Morphometrics

Leaf dimensions of *D. a.* ssp. *setchellii* are more similar to other subspecies of *D. abramsii* than to *D. cymosa* based on measurements of common garden grown plants (Table S6). In particular, *D. a.* ssp. *setchellii* (n=34) were not significantly different from either *D. cymosa* (n=18), nor other *D. abramsii* subspecies from San Luis Obispo County (n=6) for leaf length (Fig. 6A, Table S7), yet were significantly different from *D. cymosa* for maximum width (at the widest point, Fig. 6B) and for the ratio of leaf length/maximum width (Fig. 6C). *Dudleya abramsii* ssp. *setchellii* and *D. cymosa* form non-overlapping clusters in the length x maximum width plot (Fig. 6D), whereas *D. a.* ssp. *setchellii* and other subspecies of *D. abramsii* have largely overlapping length/width ratios (Tables S6 and S7).

**FIG. 6.**
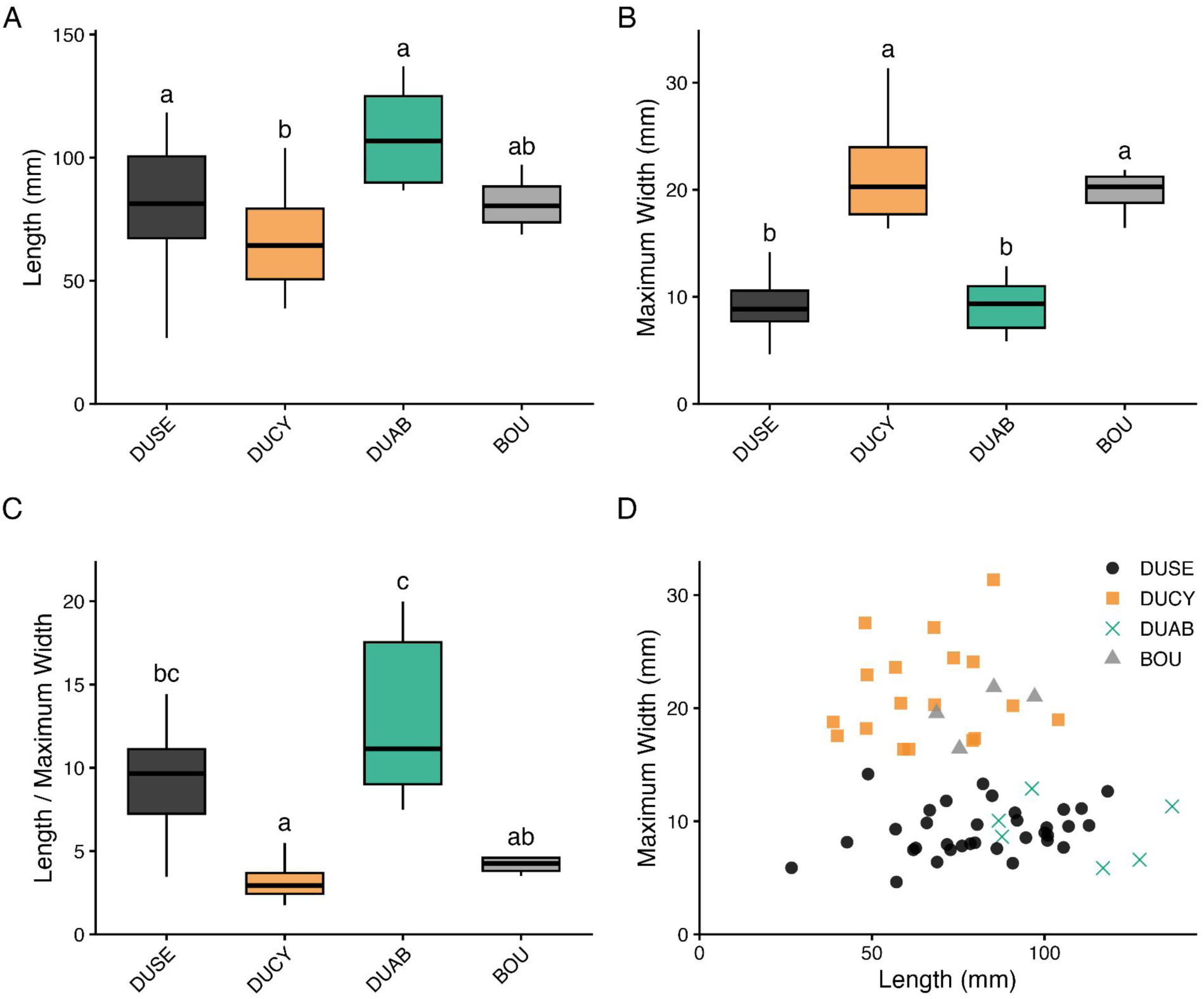
Leaf morphology of *Dudleya abramsii* ssp. *setchellii* and close relatives measured in a common garden. Box-and-whisker plots showing (A) leaf length, (B) maximum leaf width, (C) leaf shape (length / maximum width), and (D) the relationship between leaf length and maximum width. Letters indicate groupings from Holm-adjusted Dunn post hoc comparisons following significant Kruskal–Wallis tests. DUSE = *D. abramsii* ssp. *setchellii* (n = 34), DUCY = *D. cymosa* (n = 18), DUAB = two other subspecies of *D. abramsii* from San Luis Obispo County (n = 6), and BOU = Boulder Ridge population (n = 4).

We have four individuals from Boulder Ridge, the population from which CY_BOU was sampled (n=4). The four plants from the Boulder Ridge population had leaf lengths intermediate between those of *D. a.* ssp. *setchellii* and *D. cymosa* (Fig. 6A), whereas leaf widths were more similar to *D. cymosa* (Fig. 6B). The Boulder Ridge group was not significantly different from either taxon in length-to-maximum-width ratio.

### Pollinator Observations

In 2024, over two days at two sites per day, we collected ∼58 hours of camera footage with an average of 13 flowers per camera (4-41 flowers) representing ∼732 flower hours (Table S2). Overall, the bumblebee, *Bombus vosnesenskii,* was the most common pollinator of *D. a.* ssp. *setchellii* (Fig. 7). We recorded 1,956 visits by *B. vosnesenskii* (81% of all visits recorded), 333 visits by a sweat bee (Halictidae) (14%), and 120 visits from two unknown pollinators (5%). Combining the number of visits with the amount of flower hours indicates 2.67 visits/fl/h for *B. vosnesenskii*. If each flower is open for at least 2 days (Whittall unpublished observations) and able to be pollinated for a minimum of 6 h per day (12 h of availability to pollinators per flower because of climatic variables such as cool night and morning temperatures, afternoon winds, etc.), then each flower would have been visited by *B. vosnesenskii* an average of 32 times. The average *B. vosnesenskii* visit duration was 29 s compared to the longer average visit duration by sweat bees (1 min 5 s) and unknown bees (1 min 20 s).

**FIG. 7.**
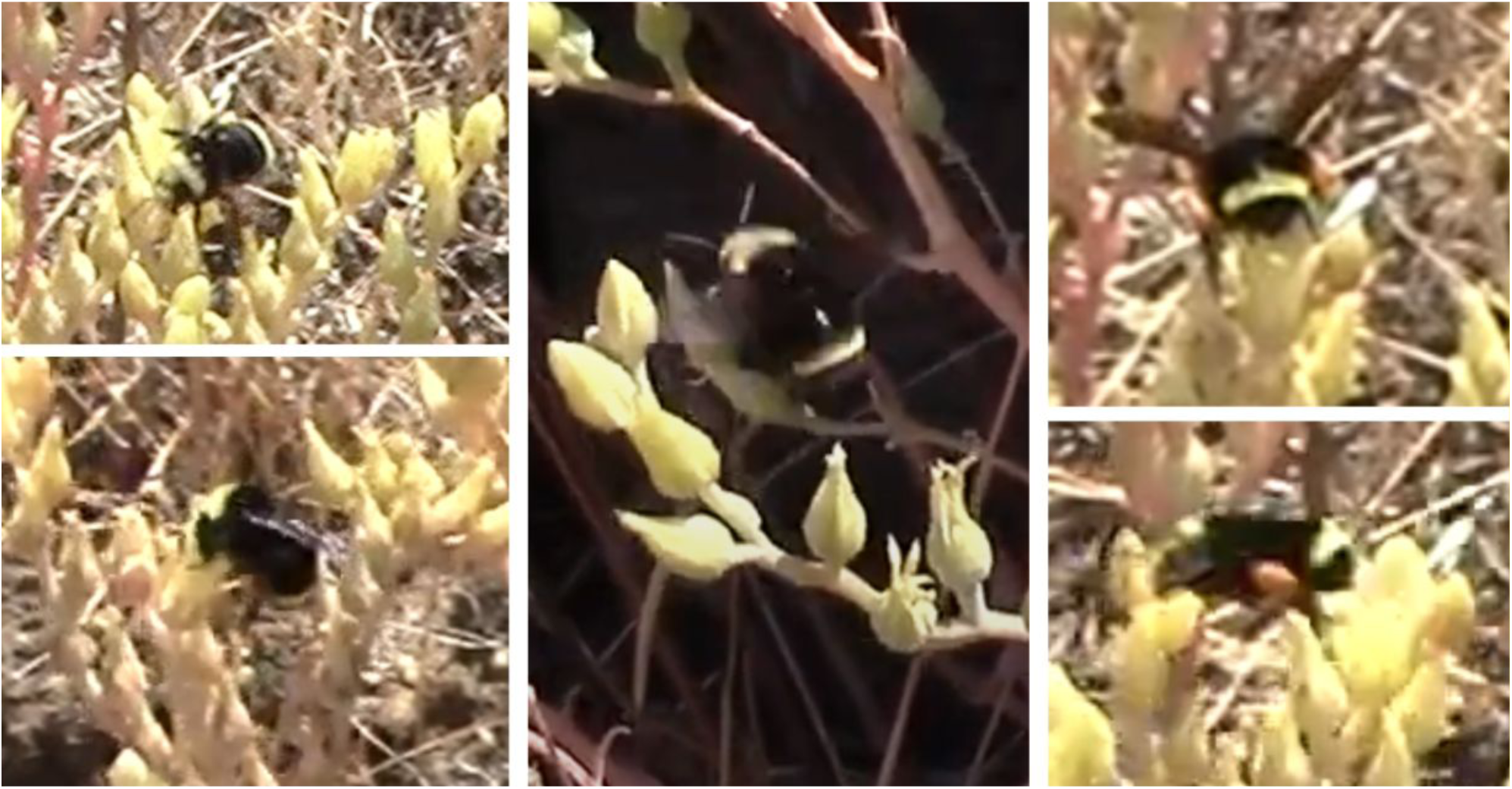
Still images extracted from pollinator recordings showing the bumblebee, *Bombus vosnesenskii*, the most common pollinator of *Dudleya abramsii* ssp. *setchellii* in spring 2024. For scale, this bumblebee species measures approximately 15 mm long.

**FIG. 8.**
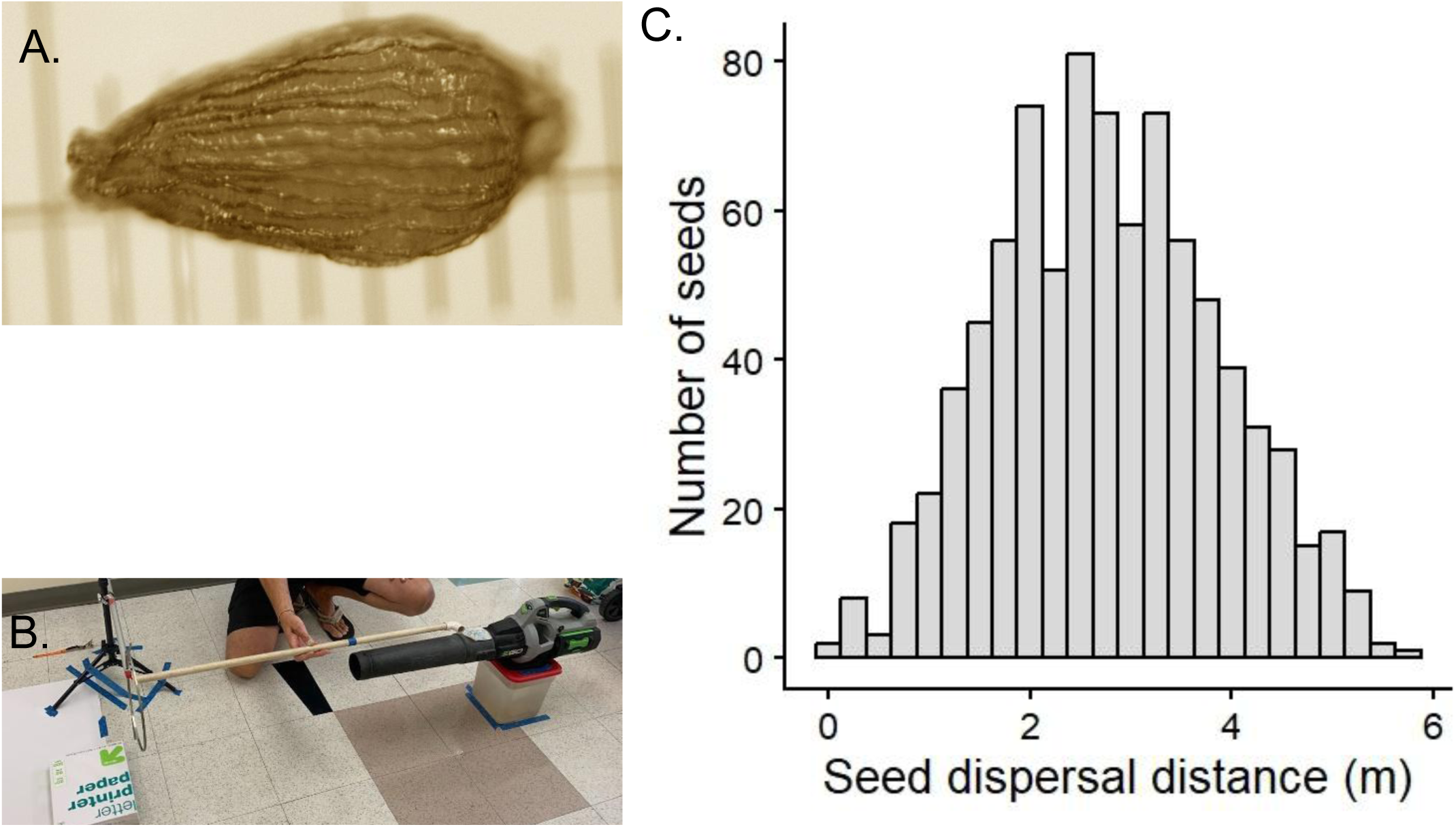
Experiment to investigate *Dudleya abramsii* ssp. *setchellii* seed characteristics and dispersal. (A) A single seed under 75× magnification with ruler markers every 0.1 mm. (B) The controlled wind dispersal experiment setup using a leaf blower. (C) Distance seeds traveled (n = 847 seeds).

### Seed Dispersal

Seeds measured approximately 0.7 × 0.4 mm, were narrowly ovoid with longitudinal ridges (striate) as described previously (McCabe 2024) - lacking any obvious structures associated with specialized long-distance dispersal (Fig. 8A). Mean dispersal distance in our artificial wind experiment was 276 cm (range: 18.3–580 cm), and the observed distances formed an approximately unimodal distribution (Fig. 8C). In total, 847 of the approximately 1,000 seeds were recovered (10 trials of 100 seeds/trial).

## Discussion

### Isolation by Distance

Isolation by distance is one of the most widely observed patterns in population genetics and represents the null expectation against which more complex hypotheses of population structure are evaluated (Wright 1943; Slatkin 1993). IBD emerges from the combined effects of limited dispersal and genetic drift. Consistent with its generality, Jenkins et al. (2010) found significant isolation by distance in approximately 60% of 240 empirical population genetic datasets spanning diverse plant and animal taxa, although the relationship typically explained only a modest proportion of genetic variation (mean r² = 0.22). Our results closely follow Wright’s (1943) original prediction, revealing significant isolation by distance both within the focal Quicksilver North population and among populations distributed throughout the range of *D. a.* ssp. *setchellii*. The strength of the within-population relationship (Mantel r² = 0.24) was remarkably similar to the cross-study average reported by Jenkins et al. (2010), whereas the among-population relationship (Mantel r² = 0.43) was nearly twice as strong, suggesting particularly pronounced geographic structuring across the naturally fragmented range of *D. a.* ssp*. setchellii*. Although these comparisons should be interpreted cautiously because Jenkins et al. (2010) synthesized diverse taxa across a broad range of spatial and temporal scales, they indicate that regional genetic differentiation in *D. a.* ssp. *setchellii* is stronger than typically observed in natural populations. In contrast to widespread species such as *Mimulus guttatus*, where extensive gene flow can be maintained across continental scales (Twyford et al. 2020), *D. a.* ssp. *setchellii* exhibits pronounced isolation by distance over remarkably fine spatial scales, reflecting its naturally fragmented distribution on serpentine habitat islands.

Our ecological observations provide insight into the dispersal component of Wright’s (1943) model. Levin and Kerster (1974) recognized that pollen generally disperses much farther than seeds in flowering plants and emphasized that these two processes contribute differently to patterns of gene flow. Building upon this framework, Epperson’s modeling study (2007) demonstrated that restricted seed dispersal often exerts a stronger influence on fine-scale spatial genetic structure because neighborhood size reflects the combined dispersal of maternal and paternal gametes. When seeds disperse primarily near maternal plants, they can create localized clusters of related individuals that may persist despite pollen-mediated gene flow. Consequently, strong isolation by distance can develop even in species with relatively extensive pollen dispersal. This interpretation is supported by studies of red oak (*Quercus rubra*), where pollen disperses substantially farther than seeds (due to anemophily), yet spatial genetic structure remains largely determined by localized seed dispersal (Moran and Clark 2012). Our results are remarkably consistent with this framework. Bumblebees (*Bombus vosnesenskii*), the dominant floral visitors of *D. a.* ssp. *setchellii* in 2024, are capable of foraging over distances considerably greater than those observed in our seed dispersal experiment, although effective pollen-mediated gene flow was not measured. For example, *B. vosnesenskii* in montane meadows has been reported to travel 1,867 m (Mola et al. 2020) and in agricultural settings, worker foraging distance was estimated to be at least 11.6 km (Rao and Strange 2012). In our indoor seed dispersal experiment, recovered seeds traveled no farther than 5.8 m under an initial airflow of 33.5 mph. Although we searched carefully beyond this distance, approximately 15% of seeds were not recovered, potentially because they escaped laterally beyond the collection surface (unrelated to their dispersal distance). However, the experiment did not reproduce natural topography, vegetation, or variable wind conditions, and therefore cannot exclude occasional longer-distance dispersal in the field. Together, these observations are consistent with limited seed dispersal contributing to the observed spatial genetic structure, although the relative contributions of seed dispersal, pollen-mediated gene flow, and genetic drift cannot be distinguished with our data.

Genetic drift, the second component of Wright’s (1943) model, may also contribute substantially to the observed genomic patterns, particularly given the small, geographically isolated populations characteristic of this serpentine endemic. Unlike widespread species occupying continuous habitat, *D. a.* ssp. *setchellii* is restricted to naturally fragmented serpentine habitat islands that support relatively small populations separated by unsuitable habitat. These populations have likely remained relatively small throughout their evolutionary history because suitable serpentine rocky outcrop habitat itself is naturally patchy. Under these conditions, repeated genetic bottlenecks and restricted dispersal would potentially reinforce one another through time, producing progressively greater genetic differentiation with increasing geographic distance (Wright 1943).

The modest increases in Mantel correlations following log transformation are consistent with genetic differentiation accumulating somewhat more rapidly over shorter geographic distances, although the differences between transformations were small, a pattern consistent with localized dispersal and strong fine-scale spatial genetic structure.

Our analyses intentionally evaluated the simplest expectation of Wright’s isolation-by-distance model by relating genetic differentiation to straight-line (Euclidean) geographic distance. More recently, landscape genetics has expanded this framework by explicitly quantifying the effects of landscape composition, habitat configuration, and matrix quality on gene flow and spatial genetic variation (Manel et al. 2003; Storfer et al. 2007).

Although *D. a.* ssp. *setchellii* occupies a relatively narrow geographic range and occurs almost exclusively on serpentine grasslands, environmental differences among sites may contribute to genetic differentiation. Further investigations of soil chemistry, aspect, elevation, vegetation, and disturbance history would help evaluate isolation by environment (Sexton et al. 2014). Nevertheless, the apparent east–west differentiation visible in the PCA, phylogenetic trees and pairwise comparisons suggests that landscape features such as Coyote Valley (present day Highway 101 cooridoor) may further restrict gene flow beyond that expected from geographic distance alone. Integrating genomic data with resistance-based landscape genetic analyses therefore represents a logical next step for distinguishing the relative contributions of geographic distance and contemporary landscape barriers to population connectivity in this endangered serpentine endemic.

Limited seed dispersal may contribute to the observed isolation by distance, but the relative contributions of seed and pollen dispersal, genetic drift, and geographic isolation remain unresolved. Nevertheless, the apparent east–west differentiation visible in the PCA, phylogenetic trees, and pairwise comparisons suggests that landscape features such as Coyote Valley may further restrict gene flow beyond that expected from geographic distance alone. Integrating genomic data with resistance-based landscape genetic analyses therefore represents a logical next step for distinguishing the relative contributions of geographic distance, dispersal limitation, and landscape barriers to population connectivity in this endangered serpentine endemic.

### Evolutionary Relationships and Hybridization

The genome-wide relationships recovered here provide new insight into the evolutionary history of *D. a.* ssp. *setchellii* while also highlighting questions that require broader taxonomic sampling. Previous phylogenetic analyses based on nuclear ITS and chloroplast DNA sequences placed *D. a.* ssp. *setchellii* within a clade containing *D. cymosa* ssp. *paniculata* and suggested a close relationship among these taxa, while also noting the potential for gene flow where distributions overlap (Yost et al. 2013). Our whole-genome resequencing results are broadly consistent with those findings but provide substantially greater resolution. Both principal component and phylogenetic analyses generally support the genomic distinctiveness of *D. a.* ssp. *setchellii*, although the identification of a putative *D. cymosa* x *D. abramsii* ssp. *setchellii* hybrid (CY_BOU) and placement of the Hunting Hollow sample indicate that relationships with *D. cymosa* require broader sampling and explicit tests of introgression. Furthermore, genomic distances indicate that *D. a.* ssp. *setchellii* is significantly more similar to nearby *D. cymosa* populations than it is to the currently recognized *D. abramsii* subspecies sampled from the closest populations in San Luis Obispo County. These results support previous suggestions that *D. a.* ssp. *setchellii* shares a relatively recent evolutionary history with *D. cymosa* (Yost et al. 2013), despite its current taxonomic placement within *D. abramsii*.

Our analyses provide evidence consistent with natural hybridization between *D. a.* ssp. *setchellii* and *D. cymosa*. CY_BOU occupied an intermediate position in principal component space and had similar mean genetic distances to the two taxa while greenhouse grown plants from the same population have intermediate leaf morphology. These combined observations support its interpretation as a putative hybrid, although explicit ancestry analyses and broader population sampling are needed to test this hypothesis. Natural hybridization has long been recognized throughout *Dudleya*, where reproductive barriers appear to be relatively weak and numerous naturally occurring hybrid combinations have been described (Moran 1951; Uhl and Moran 1953; Yost et al. 2013; Guilliams and Hasenstab-Lehman 2021). Our results therefore fit the previously reported pattern within the genus. Nevertheless, only one putative hybrid was detected despite sampling several populations, suggesting that hybridization may be relatively uncommon within *D. a.* ssp. *setchellii*.

Although rare hybridization may contribute to evolutionary novelty, it also presents a potential conservation concern. Because *D. cymosa* is considerably more widespread and abundant throughout the region than the federally endangered *D. a.* ssp. *setchellii*, repeated backcrossing to *D. a.* ssp. *setchellii* could theoretically result in asymmetrical introgression, whereby alleles from the common species gradually replace those of the rare endemic. Such genetic swamping has been recognized as an important conservation risk for rare plant species that coexist with widespread congeners (Levin et al. 1996; Rhymer and Simberloff 1996; Ellstrand and Elam 1993). Whether this process is occurring in *D. a.* ssp. *setchellii* remains unknown; however, our discovery of a putative hybrid demonstrates that opportunities for gene flow exist where these two taxa occur in close proximity. Expanded genomic sampling of additional *D. cymosa* populations, especially near zones of sympatry, together with explicit population genomic tests for introgression combined with artificial hybridization experiments will be necessary to determine the frequency, direction, and evolutionary consequences of interspecific gene flow.

There is an apparent association between genomic divergence and ecological specialization since throughout Santa Clara County, *D. a.* ssp. *setchellii* appears restricted to serpentine outcrops, whereas nearby populations of *D. cymosa* occupy predominantly non-serpentine rocky substrates despite often occurring only a few hundred meters away. Relatively few *Dudleya* taxa have been reported as serpentine-associated or serpentine-restricted (McCabe 2024; Safford and Miller 2020). These include some populations of *D. cymosa* ssp. *cymosa* north of San Francisco (Lake Co.) and in the Sierra Nevada foothills (S. McCabe, unpublished data), as well as *D. blochmaniae* ssp. *blochmaniae*. The latter is morphologically distinct from *D. a.* ssp. *setchellii* and was placed in a different major infrageneric clade by Yost et al. (2013). The persistence of a genetically cohesive serpentine endemic despite opportunities for hybridization suggests that ecological adaptation may contribute to maintaining species boundaries. Future studies integrating reciprocal transplant experiments, genome scans for adaptive loci, and broader geographic sampling may reveal whether adaptation to serpentine soils has played a central role in the origin and maintenance of *D. a.* ssp. *setchellii*.

Our common-garden comparisons also suggest an interesting disconnect between genomic affinity and morphology. Leaf width clearly separated *D. cymosa* from *D. a.* ssp. *setchellii*, whereas the other two sampled subspecies of *D. abramsii* exhibited substantial overlap with *D. a.* ssp. *setchellii*. Thus, although genome-wide relationships indicate closer affinity between *D. a.* ssp. *setchellii* and *D. cymosa*, vegetative morphology remains more consistent with its current placement alongside the other subspecies of *D. abramsii* sampled from San Luis Obispo County. This discordance illustrates why taxonomic inference based on morphology alone has proven particularly challenging within *Dudleya*. At present, our sampling is insufficient to justify formal taxonomic revision. Instead, our results provide a genomic framework that identifies several hypotheses for future testing through expanded sampling of *D. a.* ssp. *setchellii*, *D. cymosa*, and other subspecies of *D. abramsii* across their geographic ranges.

### Conservation Implications

Our results have several implications for the conservation and management of *D. a.* ssp. *setchellii* that should be considered when its conservation status is reviewed next (United States Fish and Wildlife Service 2021). First, the strong isolation-by-distance pattern detected within and among populations indicates that geographically separated populations are expected to harbor increasingly distinct genetic variation. Consequently, conserving populations throughout the species’ range will preserve substantially more evolutionary diversity than focusing conservation efforts on only a few large populations. This finding may be particularly relevant for peripheral populations, such as Communication Hill that sits at the northern edge of Coyote Ridge, which are among the most threatened, yet may also contain unique genetic variation not represented elsewhere.

Second, although clonal reproduction does occur in *D. a.* ssp. *setchellii*, it appears to be restricted to very local spatial scales in our sampling of this population. All putative clones identified in this study occurred within 8 cm of one another, suggesting that clonality is unlikely to substantially influence patterns of genetic diversity beyond the immediate neighborhood of a rosette. Nevertheless, among the 11 neighboring rosette pairs sampled within 10 cm of one another, only five (45.5%) were classified as putative clones. Thus, close physical proximity alone was not a reliable indicator of clonality. This finding has practical implications for conservation monitoring, where census size is often estimated by counting rosettes. Rosette-based counts may therefore be biased in either direction depending on the census rule used. Counting every rosette as a separate individual may overestimate the number of unique genets, whereas treating all touching rosettes as a single individual may combine genetically distinct neighbors and underestimate census size. Replicated fine-scale genomic sampling across populations could relate inter-rosette distance to the probability of clonality and support development of a standardized census protocol.

Finally, our genomic analyses tentatively support *D. a.* ssp. *setchellii* as a distinct evolutionary lineage while also demonstrating that it is genetically more similar to geographically adjacent *D. cymosa* than it is to the currently recognized *D. abramsii* subspecies sampled from San Luis Obispo County. Although these data are insufficient to justify a formal taxonomic revision, they suggest that the evolutionary relationships within this complex warrant broader genomic sampling and tests for introgression. Taken together, our findings provide a genomic framework for conserving genetic diversity across the range of this serpentine endemic while highlighting several priorities for future research.

## Conclusion

For *D. abramsii* ssp. *setchellii*, strong geographic structure, highly localized clonality in one population, and evidence of limited seed dispersal under experimental conditions emphasize the importance of conserving populations throughout the taxon’s restricted range. More broadly, our results illustrate how combining whole-genome resequencing with ecological observations can provide insights into the processes shaping genetic structure in naturally fragmented plant species and an evolutionary framework for conservation management.

## Code Availability

Code and information about the genomics bioinformatics pipeline can be found in the following GitHub repository: https://github.com/evanhackstadt/dudleya/tree/main. Data and R scripts used for the isolation-by-distance analyses, Mantel tests, and leaf-morphology analyses are also available in this repository.

## Acknowledgments

The authors acknowledge the generous funding from the US Fish & Wildlife Grant (JW), a grant from the Bureau of Reclamation’s Central Valley Project Conservation Program (JW), the SCU HPC Undergraduate Research Award (KM and EH), SCU REAL awards (DC, KM, and AS), and the DeNardo Lab Rotations (LEAP) (KM). Thanks to Daryn Baker for consistent lab support. Some of the population genomic data was contributed by the SCU students of the Winter 2026 BIOL 173 (Evolution) class who were trained by senior author (JBW) and NEB scientist coauthors (AH & SH). Thanks to the FGL and GSL staff at UC Berkeley for advice on library preparation and sequencing strategies (QB3 Genomics, UC Berkeley, Berkeley, CA, RRID:SCR_022170). Three anonymous reviewers provided thorough and thoughtful constructive criticism and suggestions for improvement on an earlier draft.

---

**Supplemental Figure S1.**
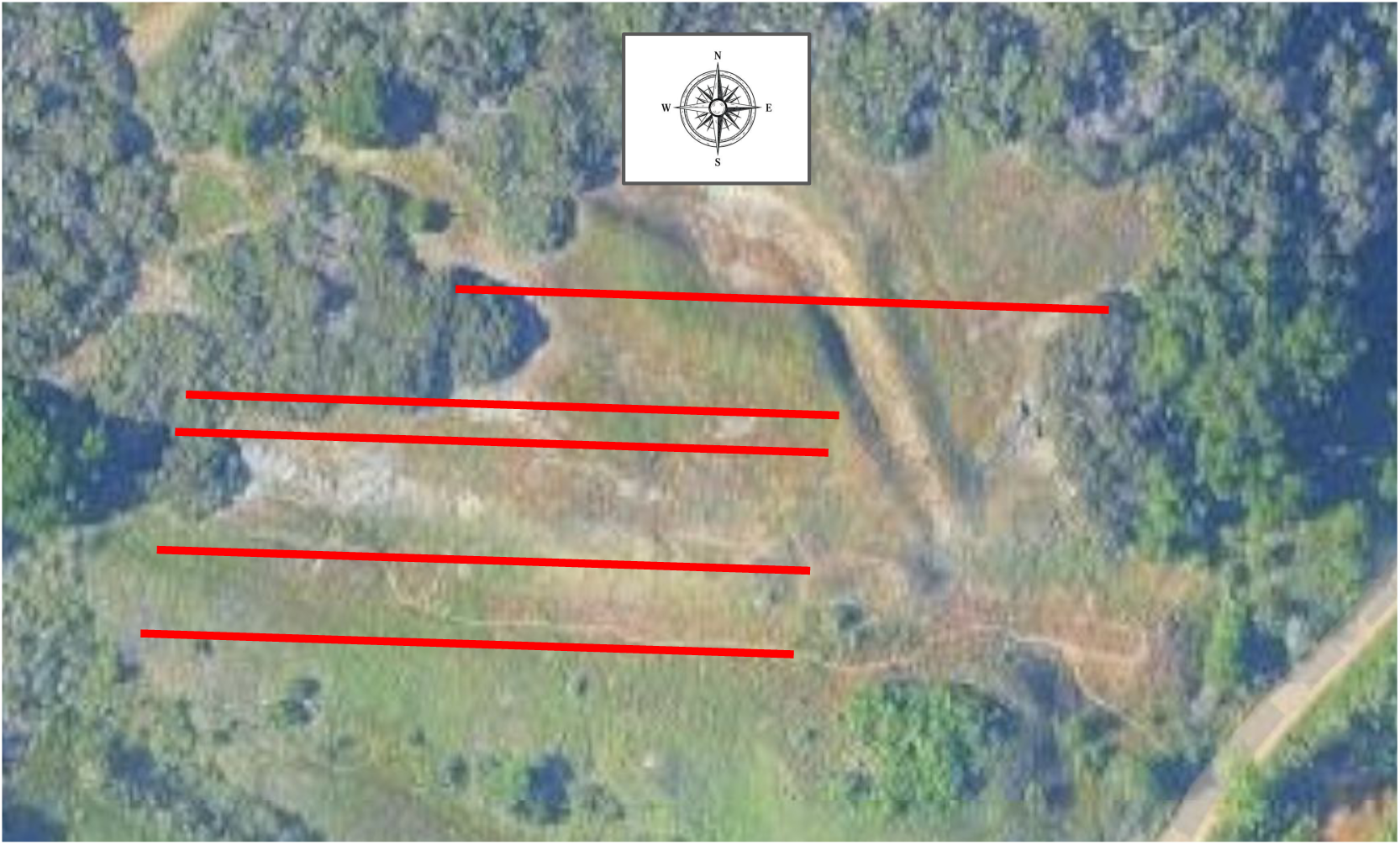
Diagram of five approximately 50–100 m transects. Lines are schematic and not drawn to scale. Plants were sampled at increasing distances from east to west to assess clonality and isolation by distance within a population.

**Supplemental Figure S2.**
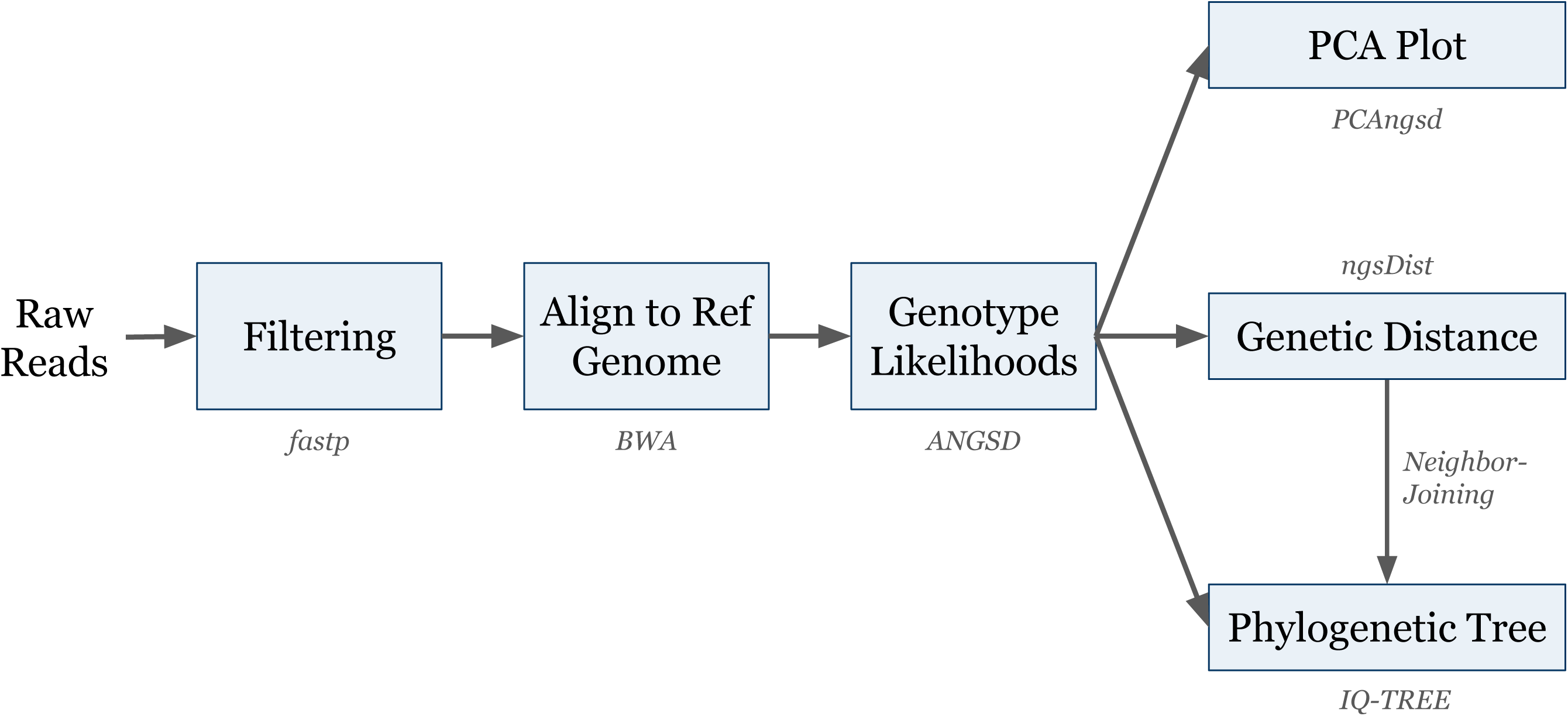
Bioinformatics workflow. Genetic distances were used to assess clonality, isolation by distance and draw a neighbor joining phylogenetic tree. Software involved is indicated in grey italicized font below the steps.

**Supplemental Figure S3.**
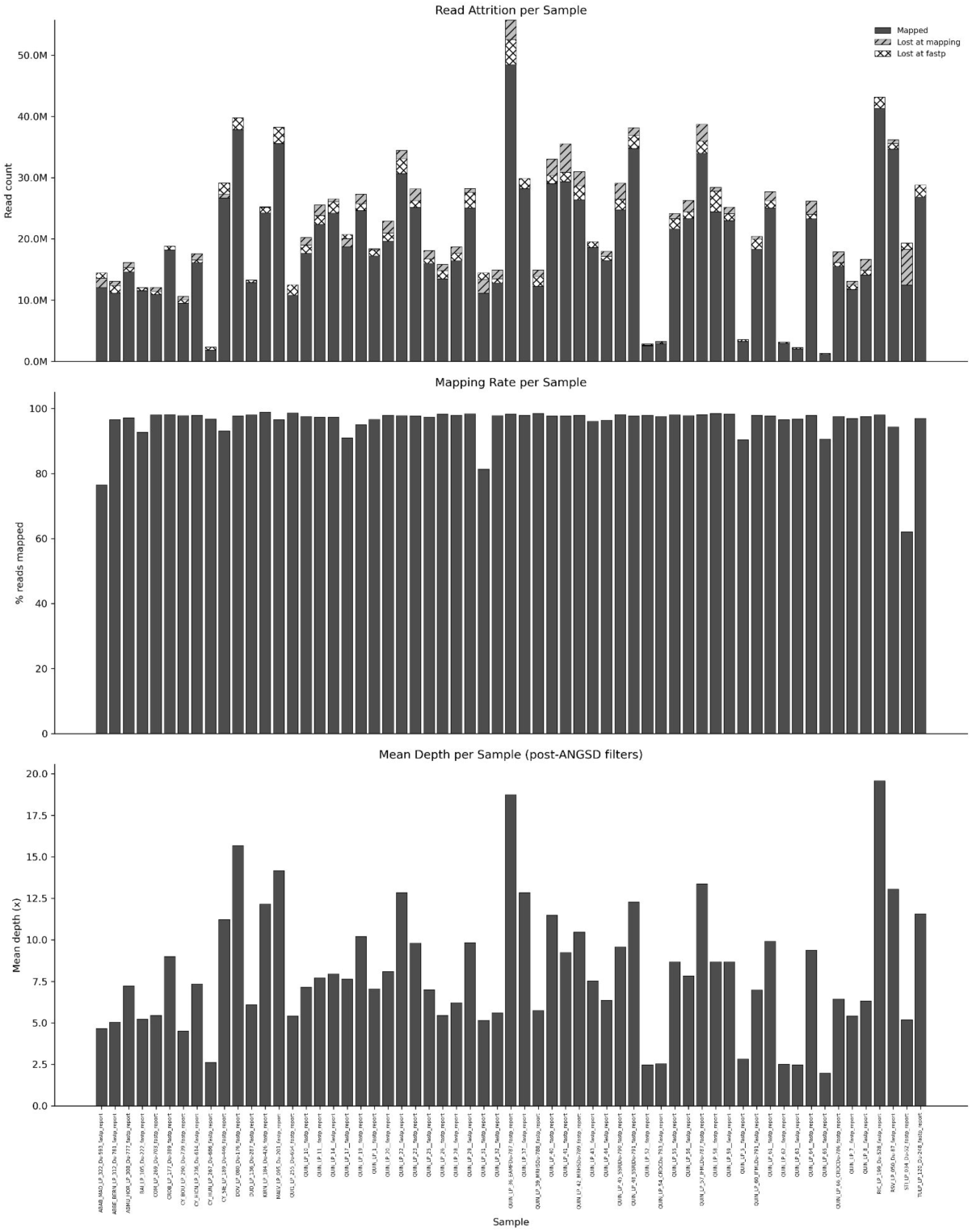
Genome resequencing results per sample indicating how many reads were preserved vs. filtered by fastp or mapping (A), percentage of reads mapped to the reference genome (B), and coverage as mean depth (C).

**Supplemental Figure S4.**
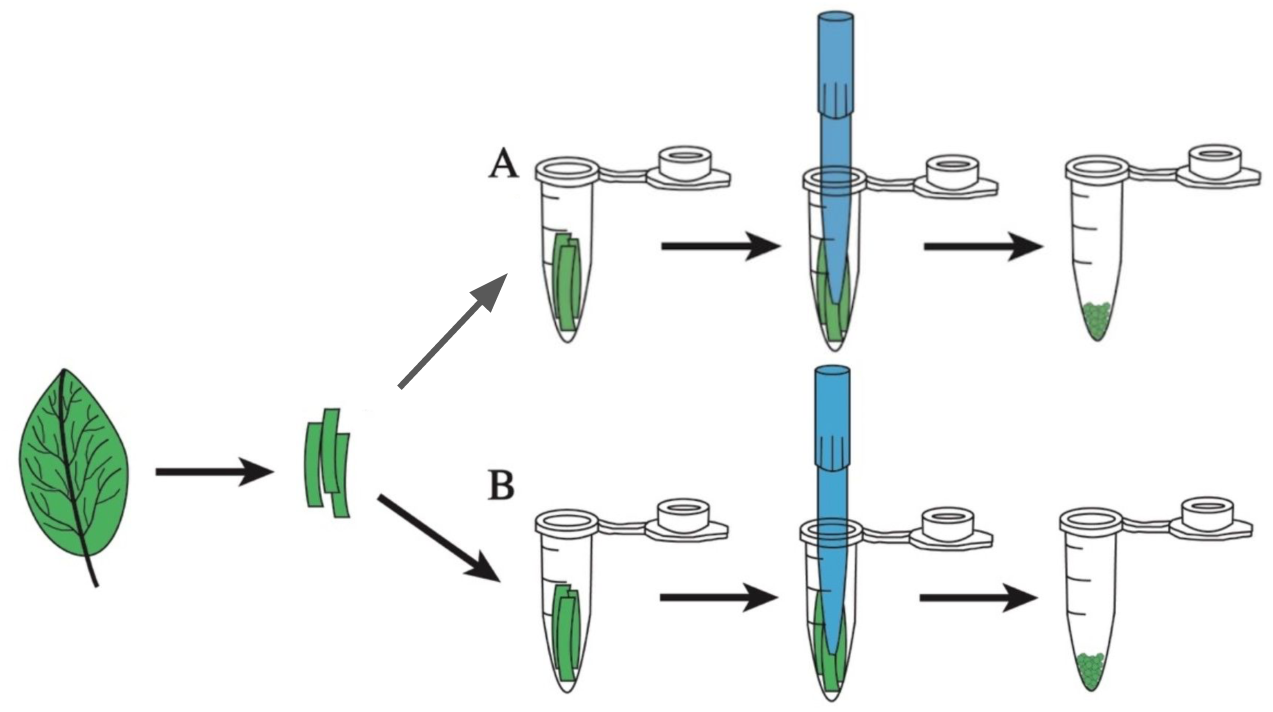
Schematic clarifying how artificial clone samples were created from a single plant, then divided into separate DNA extractions, library preps and sequencing in order to determine a threshold for putative clone assignment.

**Supplemental Figure S5.**
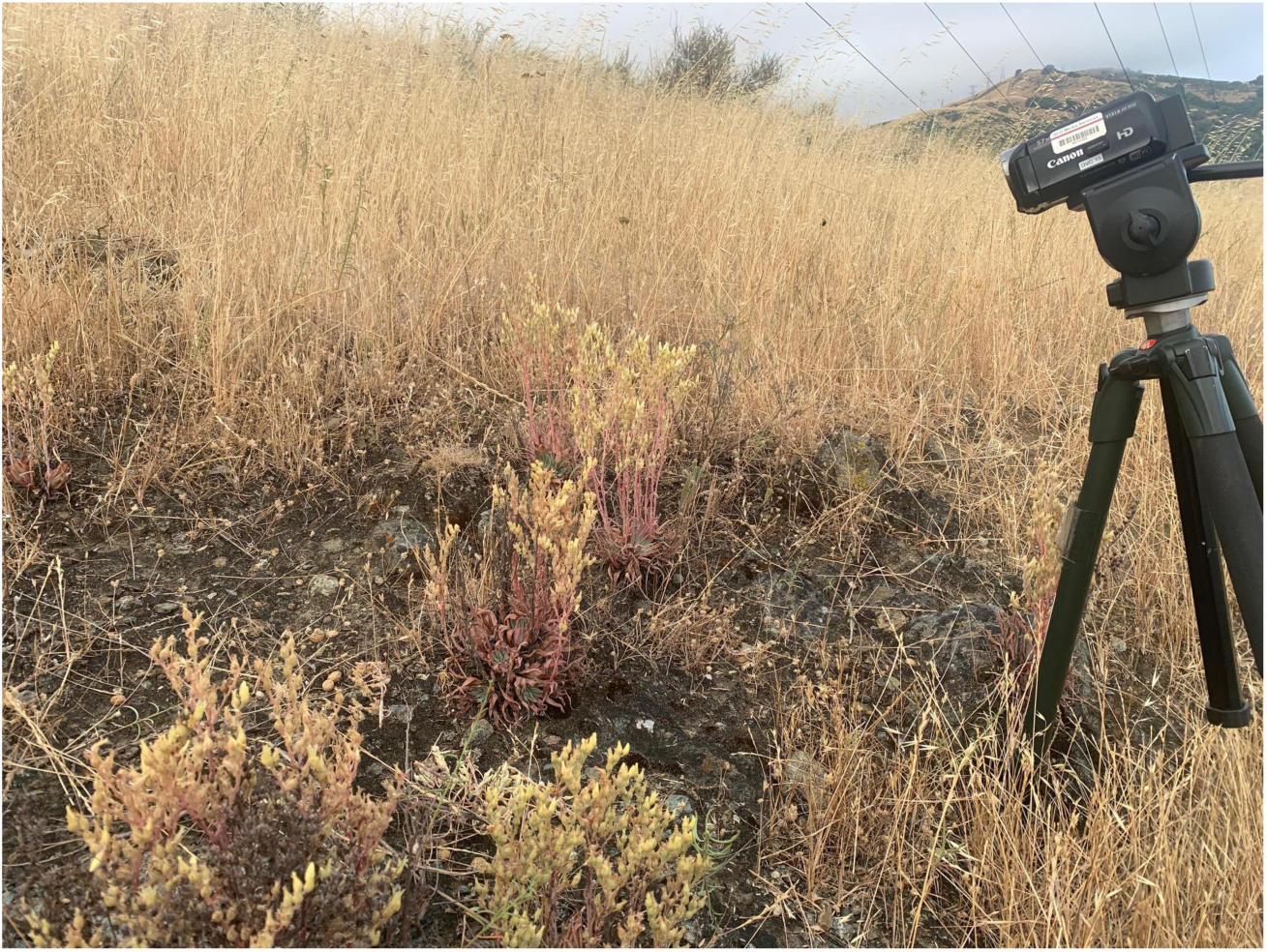
Pollinator observations were conducted using digital video camcorders as seen here at Tulare Hill. For scale, the camcorder unit is 8.5 cm long.

**Supplemental Figure S6.**
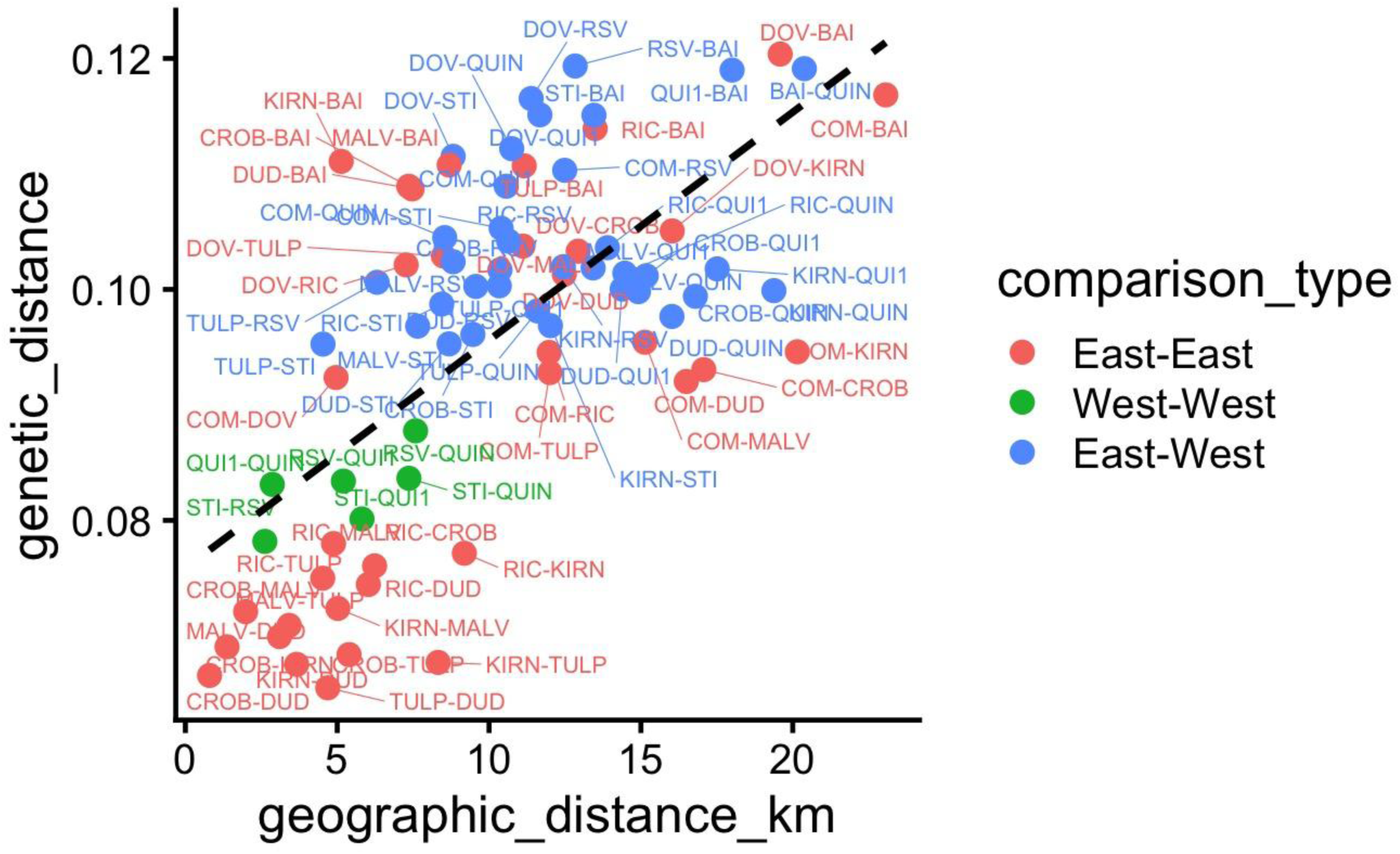
Isolation by distance plot among populations with each comparison labeled and the major groupings color coded. West includes Stiles Ranch (STI), Rancho San Vicente (RSV), Quicksilver (QUI1) and Quicksilver North (QUIN). The remaining *D. a* ssp. *setchellii* populations sampled are designated “East”.

**Supplemental Figure S7.**
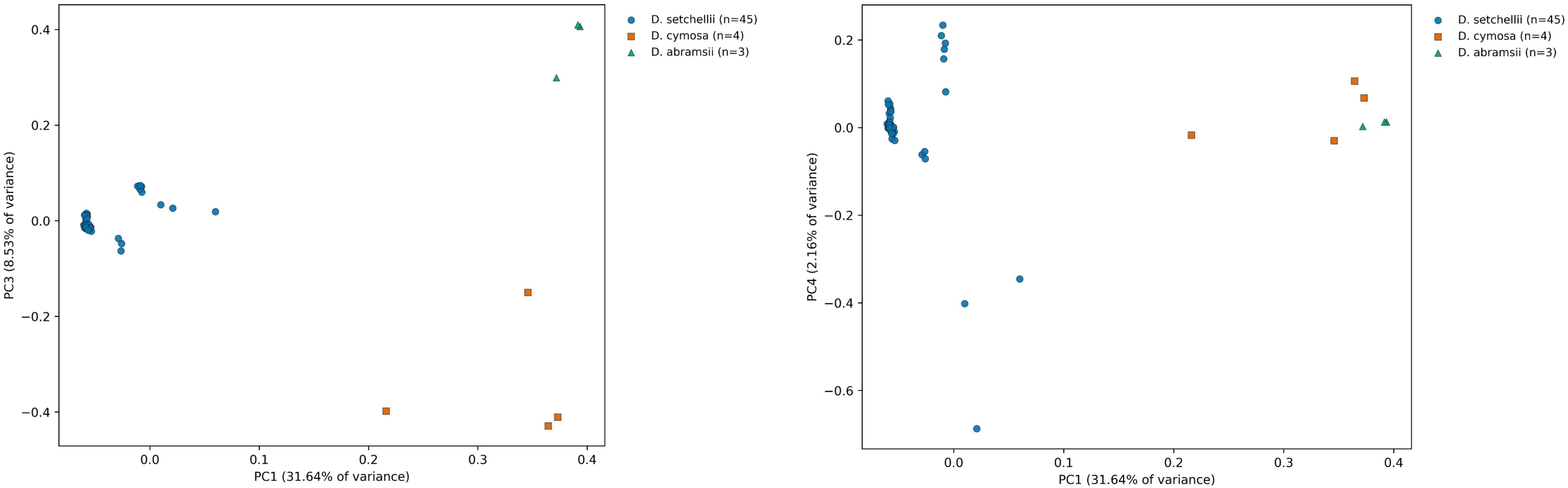
PC3 (8.53%) and PC4 (2.16%) are plotted against PC1.

**Supplemental Figure S8.**
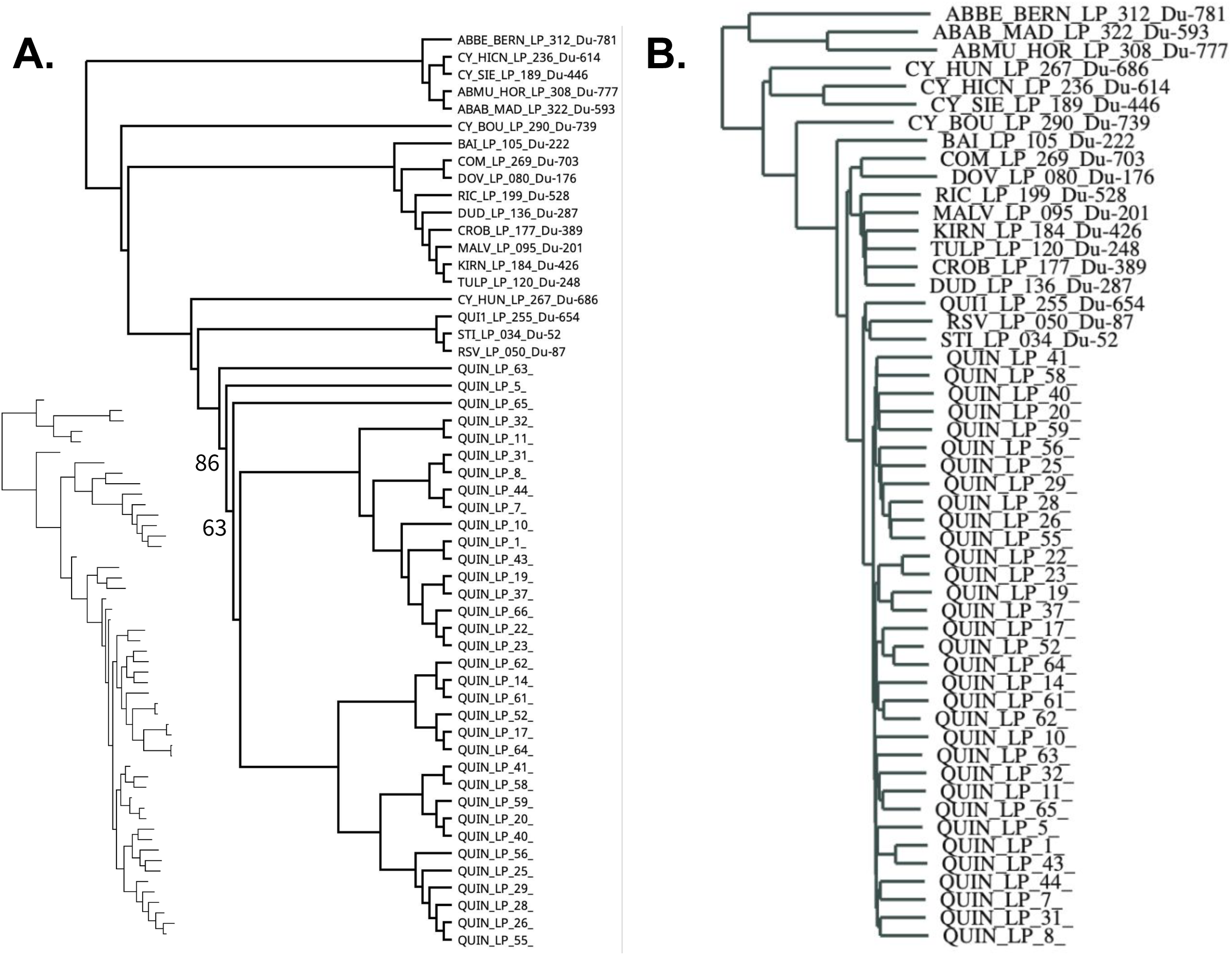
Alternative phylogenetic analysis conducted in IQTREE running maximum likelihood with GTR+ASC with 1000 bootstrap replicates (A) compared to neighbor joining phylogram with tip labels (B). Inset in (A) shows maximum likelihood phylogram with no tip labels (same order as in cladogram). In (A) all branches have 100% bootstrap except the two nodes indicated (86% and 63%).

