## Supplemental Tables & Materials for "GENE FLOW AND CONSERVATION GENOMICS OF *DUDLEYA ABRAMSII* SSP. *SETCHELLII*, AN ENDANGERED SUCCULENT IN SANTA CLARA COUNTY"

TABLE S1. SAMPLED SPECIES, POPULATIONS, THEIR CODES AND GPS COORDINATES. *All populations include a single individual except Almaden Quicksilver County Park (QUIN), where 40 individuals were sampled and 32 individuals were retained for final population genomic analyses. ** San Luis Obispo County, CA (all other samples are from Santa Clara County, CA).

| **Species** | **Population Name** | **Code** | **Latitude (°N)** | **Longitude (°W)** | **Elevation (m)** | **Substrate** |
|---|---|---|---|---|---|---|
| *D. a.* ssp. *setchellii* | Baird Ridgeline | BAI | 37.143716 | -121.674902 | 190 | Serpentine |
| *D. a.* ssp. *setchellii* | Communication Hill | COM | 37.286387 | -121.863931 | 117 | Serpentine |
| *D. a.* ssp. *setchellii* | CROSP Bench | CROB | 37.207058 | -121.698827 | 176 | Serpentine |
| *D. a.* ssp. *setchellii* | Dove Road | DOV | 37.284464 | -121.807903 | 87 | Serpentine |
| *D. a.* ssp. *setchellii* | Dude Rock | DUD | 37.205604 | -121.707635 | 177 | Serpentine |

| | | | | | | |
|---|---|---|---|---|---|---|
| *D. a.* ssp. *setchellii* | Kirby North | KIRN | 37.189776 | -121.671358 | 318 | Serpentine |
| *D. a.* ssp. *setchellii* | Malech Vista | MALV | 37.213127 | -121.719898 | 143 | Serpentine |
| *D. a.* ssp. *setchellii* | Almaden Quicksilver North* | QUIN* | 37.212085 | -121.888355 | 128 | Serpentine |
| *D. a.* ssp. *setchellii* | Almaden Quicksilver Tr 1 | QUI1 | 37.191447 | -121.8690406 | 311 | Serpentine |
| *D. a.* ssp. *setchellii* | Richmond Ranch | RIC | 37.253395 | -121.737163 | 291 | Serpentine |
| *D. a.* ssp. *setchellii* | Rancho San Vicente | RSV | 37.182103 | -121.811501 | 224 | Serpentine |
| *D. a.* ssp. *setchellii* | Stiles Ranch | STI | 37.205181 | -121.805689 | 176 | Serpentine |
| *D. a.* ssp. *setchellii* | Tulare Hill | TULP | 37.221648 | -121.757448 | 159 | Serpentine |
| *D. cymosa* | Boulder Ridge | CY_BOU | 37.232023 | -121.860634 | 131 | Non-serpentine |
| *D. cymosa* | Sierra Azul | CY_SIE | 37.189656 | -121.953911 | 476 | Serpentine |
| *D. cymosa* | Hicks Rd. North | CY_HICN | 37.195572 | -121.8762392 | 197 | Non-serpentine |
| *D. cymosa* | Hunting Hollow | CY_HUN | 37.075781 | -121.462337 | 278 | Non-serpentine |
| *D. abramsii* ssp. *bettinae* | San Bernardo Cr. Rd.** | DUABBE_BERN | 35.387158 | -120.773095 | 104 | Serpentine |

| | | | | | | |
|---|---|---|---|---|---|---|
| *D. abramsii* ssp. *murina* | Madonna Inn** | DUABAB_MAD | 35.265114 | -120.678501 | 86 | Serpentine |
| *D. abramsii* ssp. *murina* | Horse Canyon** | DUABMU_HOR | 35.319352 | -120.655306 | 260 | Serpentine |

TABLE S2. SUMMARY OF CAMERA RECORDING HOURS AND FLOWERS HOURS PER SITE AND PER DAY REPORTED IN HRS:MIN:SEC.

| | **Tulare** | | **Kirby** | | **Total** | |
|---|---|---|---|---|---|---|
| **Day** | **Total Camera Hrs** | **Total Flower Hrs** | **Total Camera Hrs** | **Total Flower Hrs** | **Total Camera Hrs** | **Total Flower Hrs** |
| 6/13 | 17:57:04 | 414:52:21 | 8:22:35 | 122:24:47 | 26:19:39 | 537:17:08 |
| 6/18 | 20:11:55 | 125:33:23 | 11:35:28 | 69:31:36 | 31:47:23 | 195:04:59 |
| TOTAL | 38:08:59 | 540:25:44 | 19:58:03 | 191:56:23 | 58:07:02 | 732:22:07 |

TABLE S3. WIND VELOCITIES AT VARYING DISTANCES FROM THE DROP ZONE.

| Distance from Drop Zone (meters) | Wind Velocity (mph) |
|---|---|
| 0 | 33.5 |
| 1 | 12.4 |
| 2 | 7.5 |
| 3 | 5.6 |
| 4 | 4.1 |
| 5 | 3.4 |
| 6 | 3.1 |

TABLE S4. GENOME SIZE ESTIMATION OF FIVE *D. A.* SSP. *SETCHELLII* SAMPLES FROM LIFEASIBLE.

| **Sample #** | **Genome Size (pg/2C)** |
|---|---|
| FC21 | 0.61 |
| FC22 | 0.62 |
| FC23 | 0.59 |
| FC24 | 0.58 |
| FC25 | 0.61 |
| Mean | 0.602 |

TABLE S5. ISOLATION BY DISTANCE COMPARISONS (MANTEL TEST RESULTS) COMPARING RAW GEOGRAPHIC DISTANCES VS LOG TRANSFORMED GEOGRAPHIC DISTANCES.

| | Within Population Mantel r | Among Populations Mantel r |
|---|---|---|
| Linear | 0.490 | 0.656 |
| $Log_{10}$ Transformed | 0.524 | 0.715 |
| Transformation Improvement | 0.034 | 0.059 |

TABLE S6. LEAF MORPHOLOGY MEAN VALUES.

| Species | N plants | Length (mm) | Width at Midpoint (mm) | Width at Widest (mm) | Width at Base (mm) |
|---|---|---|---|---|---|
| DUAB | 6 | 109.69 ± 11.77 | 9.58 ± 0.92 | 9.22 ± 1.11 | 9.03 ± 0.91 |
| DUSE | 34 | 82.38 ± 3.61 | 9.42 ± 0.39 | 9.15 ± 0.37 | 7.91 ± 0.30 |
| DUCY | 18 | 64.57 ± 3.38 | 18.52 ± 0.94 | 21.27 ± 1.02 | 15.71 ± 1.06 |
| BOU (putative hybrid)* | 4 | 91.50 ± 9.72 | 17.15 ± 1.20 | 19.71 ± 1.19 | 12.95 ± 1.17 |

* BOU = four plants from the Boulder Ridge population from which the genomic sample CY_BOU was collected.

TABLE S7. LEAF MORPHOLOGY STATISTICAL RESULTS.

| Trait | Kruskal_statistic | Kruskal_p |
| --- | --- | --- |
| Length (mm) | 15.2 | 0.00162 |
| Width at Midpoint (mm) | 42.0 | 4.11E-09 |
| Width at Widest | 41.9 | 4.15E-09 |
| Width at Base | 41.3 | 5.77E-09 |
| Ratio_Length_Midpoint | 41.3 | 5.70E-09 |
| Ratio_Length_Widest | 40.6 | 8.00E-09 |
| Ratio_Length_Base | 37.5 | 3.63E-08 |

Supplemental materials

“matrix2tree” is custom code for generating neighbor joining phylogenetic trees and available using this Figshare link (https://figshare.com/account/articles/34878522?file=70958040) and this Figshare DOI (10.6084/m9.figshare.34878522).

Genetic distances and geographic distance matrices and associated R script used for our IBD analyses within the QUIN population and among populations are also available in Figshare (https://figshare.com/s/fd0fc3fdbe89682475a7).
